# Asymmetrical diversification of the receptor-ligand interaction controlling self-incompatibility in Arabidopsis

**DOI:** 10.1101/734079

**Authors:** Chantreau Maxime, Céline Poux, Marc F. Lensink, Guillaume Brysbaert, Xavier Vekemans, Vincent Castric

## Abstract

How two-components genetic systems accumulate evolutionary novelty and become diversified in the course of evolution is a fundamental problem in evolutionary systems biology. In the Brassicaceae, self-incompatibility (SI) is a spectacular example of a diversified allelic series in which numerous highly diverged receptor-ligand combinations are segregating in natural populations. However, the evolutionary mechanisms by which new SI specificities arise in the first place have remained elusive. Using *in planta* ancestral protein resurrection, we demonstrate that two allelic variants currently segregating as distinct receptor-ligand combinations diverged through an asymmetrical process whereby one variant has retained the same recognition specificity as the (now extinct) ancestor, while the other has functionally diverged and now represents a novel specificity no longer recognized by the ancestor. Examination of the structural determinants of the shift in binding specificity suggests that allosteric changes may be an important source of evolutionary novelty in this highly diversified receptor-ligand system.

## Introduction

A central goal of biology is to understand the evolutionary forces and molecular processes by which new traits and functions emerge in living organisms. A major challenge toward this goal is that many cellular processes rely on interactions between molecular partners (protein-protein interactions or regulatory interactions between e.g. transcription factors and their binding sites; Boyle et al., 2017; Courtier-Orgogozo et al., 2019) rather than on the action of individual components in isolation. At present, the overall structure of protein-protein or regulatory interaction networks is just starting to be deciphered at the genome level in a handful of model organisms, and a general understanding of how the diversification of these networks led to the emergence of new biological functions in distinct lineages is crucially lacking (Andreani and Guerois 2014; Nooren and Thornton 2003). In two-components genetic systems, specificity of the interaction can be very tight such as e.g. in receptor-ligand interactions (Laub & Goulian, 2007) or bacterial toxin-antitoxin systems (Aakre et al., 2015). In these systems, the emergence of novel traits involves evolutionary modification of the interacting partners, leading to potential disruption of the interaction or to detrimental cross-talk, at least transiently (Plach et al., 2017). A central aspect of the diversification process therefore concerns the functional nature of the evolutionary intermediate. Two main scenarios have been proposed (Aakre et al., 2015). The first scenario posits that a functional change results from introduction of a mutation that first modifies one of the two partners, initially disrupting the functional interaction and leading to a non-functional intermediate. If fitness of this non-functional intermediate is not impaired too drastically, it may remain in the population until a compensatory mutation hits the second partner and rescues the interaction in a modified state, resulting in the novel function. An alternative scenario though, is that the first mutation may broaden rather than inactivate specificity of one of the two interacting partners, transiently releasing constraint on the other before specificity of the first partner becomes restricted again, thus maintaining functionality of the system all along the process. Distinguishing between these two scenarios requires functional characterization of the ancestral evolutionary intermediates, which has been achieved in a very limited number of biological cases such as bacterial toxin-antitoxin systems (Aakre et al., 2015), acquisition of cortisol specificity of the mammalian glucocorticoid receptor (Bridgham et al., 2009) or mammalian retinoic acid receptors (Gutierrez-Mazariegos et al., 2016). The scenario of a promiscuous intermediate seems to have received better support, but whether this is a truly general pattern remains to be determined. A major limitation is that detailed population genetics models taking into account how natural selection is acting on variants of the two genes given their specific biological function are lacking for most genetic systems (Aakre et al., 2015; Aharoni et al., 2005; Bloom and Arnold 2009; Bridgham et al., 2009; Matsumura and Ellington 2001; Matton et al., 2000; Sayou et al., 2014).

Self-incompatibility (SI) in the flowering plants is a prime biological system to investigate how functional diversification can proceed between two molecular partners over the course of evolution. SI has evolved as a strategy to prevent selfing and enforce outcrossing, thereby avoiding the deleterious effects of inbreeding depression (Kitashiba and Nasrallah 2014; Nettancourt 1977). In *Brassicaceae*, SI is genetically controlled by a single multiallelic non-recombining locus called the S-locus (Nettancourt 1977). This locus encodes two highly polymorphic self-recognition determinant proteins: the female cell-surface ‘receptor’ S-LOCUS RECEPTOR KINASE (SRK) is expressed in stigmatic papillae cells (Takasaki et al., 2000) and the male ‘ligand’ S-LOCUS CYSTEIN RICH PROTEIN (SCR) is displayed at the pollen surface (Schopfer et al., 1999; Takasaki et al., 2000; Takayama et al., 2000). The SI response consists in pollen rejection by the pistil and is induced by allele-specific interaction between the SRK and SCR proteins when encoded by the same haplotype (S-haplotype, Kachroo et al., 2001; Nasrallah & Nasrallah, 1993; Takayama et al., 2001). These two interacting proteins exhibit a high degree of sequence variability and a large number of these highly diverged allelic variants segregate in natural populations of SI species (several dozens to nearly two hundreds, Busch et al., 2014; Castric & Vekemans, 2004; Lawrence, 2000). This large allelic diversity of receptor-ligand combinations must have arisen through repeated diversification events, but the molecular process and evolutionary scenario involved in the diversification of an ancestral haplotype into distinct descendant specificities are still unresolved.

Three evolutionary scenarios for emergence of new self-incompatibility specificities have been proposed so far (Charlesworth et al., 2005). The “compensatory mutation” scenario posits that diversification proceeds through self-compatible intermediates i.e. following transient disruption of the receptor-ligand interaction (Fig S1A, Gervais et al., 2011; Uyenoyama et al., 2001). Population genetic analysis confirmed that S-allele diversification is indeed possible through this scenario, but only under some combinations of model parameters (high inbreeding depression, high rate of self-pollination and low number of co-segregating S-alleles, Gervais et al., 2011). Under these restrictive conditions, the ancestral recognition specificity can be maintained in the long run along with the derived specificity, effectively resulting in allelic diversification. A second scenario was presented by Chookajorn et al., (2004), who proposed instead that diversification resulted from the progressive reinforcement of SRK/SCR recognition capacity among slight functional variants of *SCR* and *SRK* that might segregate within the population. This model assumes no SC intermediate, but predicts rapid functional divergence along allelic lineages. A similar « replacement » dynamics was indeed observed by Gervais et al., (2011), whereby the introduced self-compatible intermediate excluded its functional ancestor from the population under a large portion of the parameter space, such that secondary introduction of the compensatory mutation effectively resulted in the turnover of recognition specificities along allelic lines rather than in diversification *per se*. In this model, a turnover of recognition specificities may therefore be expected along each allelic lineage, rather than their long-term maintenance over evolutionary times (Fig S1B). A third scenario was proposed by Matton et al. (1999) and involves a dual-specificity intermediate (Fig S1C). This scenario was criticized on population genetics arguments because such a dual-specificity haplotype would recognize and reject more potential mates for reproduction than its progenitor haplotype resulting in lower reproductive success, and should therefore be disfavoured by natural selection and quickly eliminated from the populations (Charlesworth 2000; Uyenoyama and Newbigin 2000). Overall, while the large allelic diversity at the S-locus is one of the most striking features of this system, the evolutionary scenario and molecular route by which new S-alleles arise remain largely unknown. An essential limitation in the field is the crucial absence of direct experimental approaches.

## Results

To conclusively contrast these models we used ancestral sequence reconstruction of S-haplotypes that are currently segregating in the plant *Arabidopsis halleri*. Ancestral sequence reconstruction is a new approach to resurrect ancestral biological systems, whereby the sequence of an ancestral gene or allele is determined through phylogenetic analyses, then synthesized *de novo* and its functional properties (in terms of *e.g*. biochemical activity, conformation or binding capacity) are compared to those of its contemporary descendants. Although most S-haplotypes in *A. halleri* are typically so highly diverged that even sequence alignment can be challenging, previous work identified a specific pair of S-haplotypes (S03 and S28) with high phylogenetic proximity based on a fragment of the *SRK* gene (Castric et al., 2008). We took this opportunity to reconstruct, resurrect and phenotypically characterize their last common ancestral receptor SRKa (“a” for ancestor, Fig 1) into *A. thaliana*, which has previously been established as a model plant for mechanistic studies of SI (Nasrallah et al., 2002; Tsuchimatsu et al., 2010).

**Figure 1:**
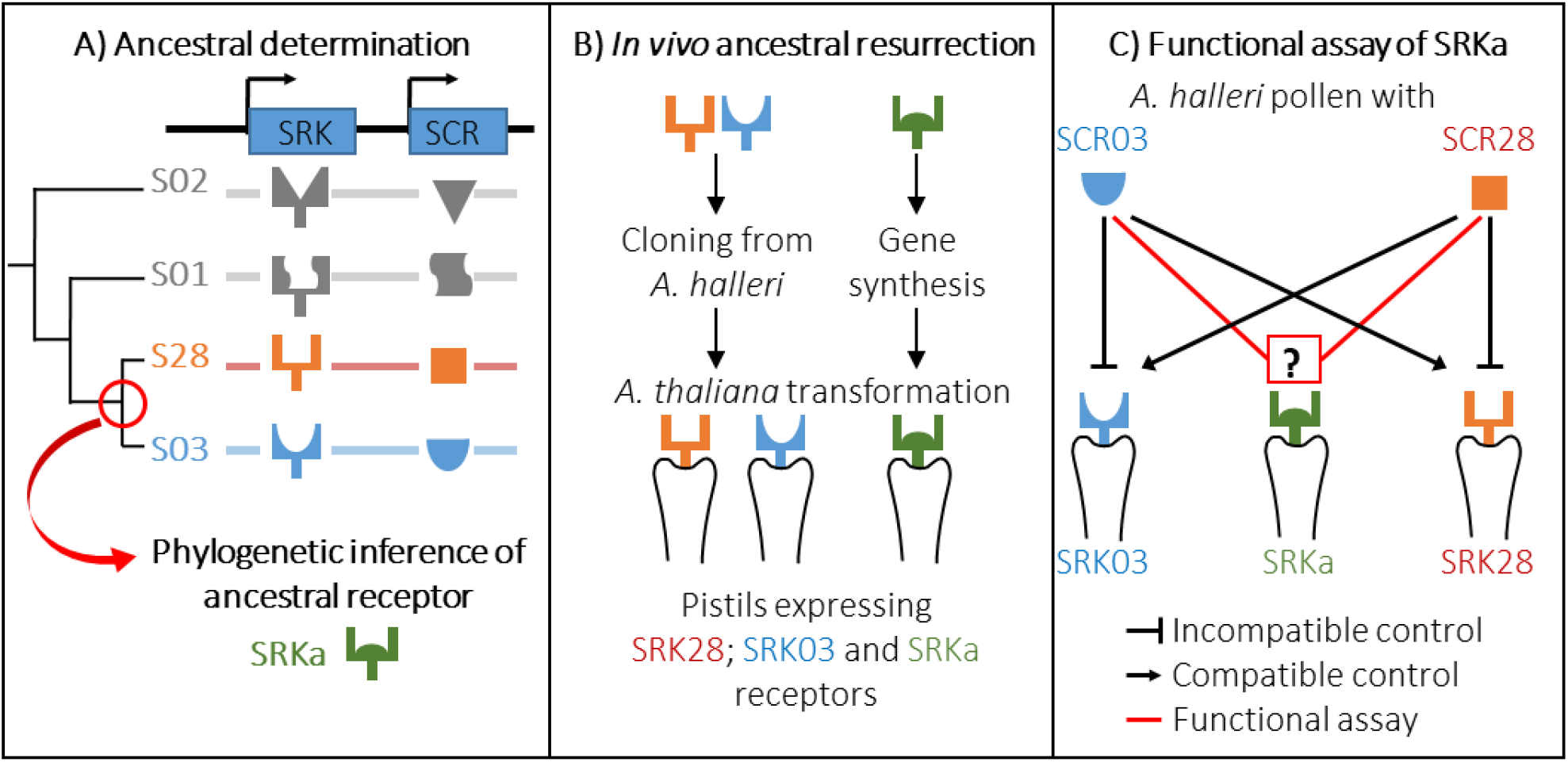
Experimental approach for the ancestral resurrection experiment. **(A)** The sequence of the common ancestor of SRK03 and SRK28 was inferred by a phylogenetic approach using codon-based models implemented in PAML. Four different versions of SRKa were defined due to inference uncertainty at two aa positions. **(B)** SRK03 and SRK28 sequences were cloned from *A. halleri* DNA BAC clones, whereas SRKa sequences were obtained by gene synthesis. **(C)** Representation of the controlled cross program to decipher the specificity of SRKa.

### Expression of S03 and S28 recognition specificities in *A. thaliana* by genetic transformation

Because of their relatively high sequence similarity, we first ensured that the S03 and S28 recognition specificities are indeed phenotypically distinct when expressed in *A. thaliana*. It was previously shown that the self-compatible model plant *A. thaliana* can mount a functional self-incompatibility response upon introduction of the S-locus genes *SCR* and *SRK* (Boggs et al., 2009; Durand et al., 2014; Nasrallah et al., 2002) but it is unclear whether transfer is possible for all *SCR* and *SRK* variants. We first assessed the activity of *AhSRK03* and *AhSRK28* promoters in transformed *A. thaliana* lines (Fig S2), which enabled us to identify the temporal window of *SRK* expression (Fig S3). Then, we used *A. halleri* pollen carrying either the S03 or S28 male determinant to validate that a proper SI response was indeed successfully transferred in transformed *A. thaliana* lines for each of our *SRK* receptors, and that this response was strictly specific toward either of their cognate ligand (two replicate transgenic lines for SRK03 and SRK28 each, Figure 2).

**Figure 2:**
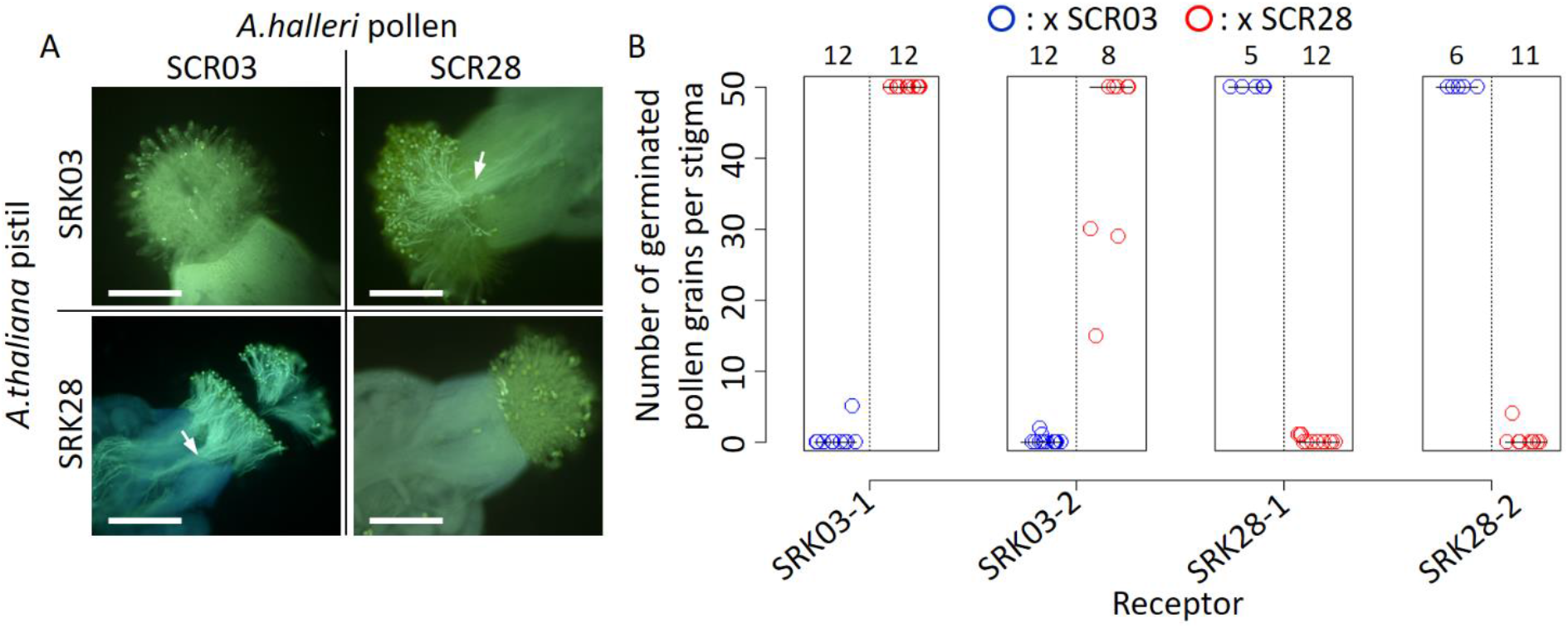
SRK03 and SRK28 are successfully expressed in A. thaliana and represent phenotypically distinct female recognition specificities. **(A)** Fluorescence microscopic observation of pistils from SRK-transformed A. thaliana plants pollinated with S03 and S28 pollen from A. halleri. A robust self-incompatibility reaction is observed between cognate alleles, whereas a compatible reaction is observed between non-cognate alleles. White arrows indicate pollen tubes that germinated in the stigma, and are specific to compatible reactions. Bar = 0.3 mm. **(B)** Number of germinated pollen grains per stigma after pollination. Two SRK03 and SRK28 lines were pollinated with A. halleri pollen expressing either S03 (blue) or S28 (red) specificities. The number of pollinated pistils for each pollination assay is indicated on the top of the figure. The median value for each cross is represented by a horizontal bar.

Abundant germination of pollen tubes occurred in crosses involving non-cognate SRK and SCR proteins, whereas crosses involving proteins from the same S-haplotype induced inhibition of pollen germination, which is characteristic of an incompatible response. Intensity of the incompatible response was then quantified by counting the number of germinated pollen grains at the pistil surface of two selected lines for each transgene (Fig 2A, 2B). Incompatible responses are characterized by less than 10 germinated pollen tubes whereas compatible crosses induced significant pollen germination with more than 50 pollen tubes per stigma. Hence, we demonstrate both that cross-species pollinations of A. *thaliana* with *A. halleri* pollen are able to induce a robust SI reaction, and also that S03 and S28 are currently fully distinct recognition specificities despite their close phylogenetic proximity.

### Phylogeny-based ancestral SRK reconstruction

We then used a phylogenetic approach to reconstruct the sequence of the most recent common ancestor of *SRK03* and *SRK28* (Fig S4). Based on a set of closely related *SRK* sequences from Arabidopsis and Capsella, we used the best fitting codon-based models implemented in PAML (Yang 2007) to reconstruct the most likely ancestral amino acid sequence (SRKa^IA^). For the vast majority of amino acid residues where *SRK03* and *SRK28* differ, the ancestral reconstruction had high posterior probability (>0.95), but was more uncertain at four positions (Fig S5). Two of these sites were close (208) or within (305) hyper variable regions known to be functionally important (Fig S4, Kusaba et al., 1997). To take this uncertainty into account and evaluate the impact of variation at these individual sites, three additional ancestral sequences were generated and tested: SRKa^IG^, SRKa^MA^ and SRKa^MG^. To avoid context-dependent effects, each of these four versions of *SRKa* was then surrounded by a promoter and a kinase sequence corresponding either to the native SRK03 (p03_SRKa_k03) or the native SRK28 (p28_SRKa_k28) variant. For each of these constructs, we obtained several replicate transgenic lines, resulting in a total of 28 different fixed homozygote single-copy transformed *Arabidopsis* lines (8 *SRKa^IA^* lines; 7 *SRKa^IG^* lines; 7 *SRKa^MA^* lines and 6 *SRKa^MG^* lines). Five lines showed very low levels of *SRKa* transcripts in pistils (Fig S6; possibly due to positional effects as is common in transgenic constructs) and were thus excluded from further analysis.

### Ligand specificity of the ancestral SRK

We then evaluated the recognition phenotype conferred by these resurrected ancestral SRK sequences by performing controlled pollination assays for all remaining 23 *SRKa* lines with pollen from *A. halleri* S03, *A. halleri* S28 and *A. thaliana* C24 (used as a control to validate receptivity of the acceptor pistil). The SI reaction was scored as the number of germinated pollen grains per stigma. All 23 tested ancestral lines showed intense pollen germination with C24 pollen, indicating that transgene integration in itself did not impair pistil fertility or ability to receive pollen (Fig 3 and Fig S7).

**Figure 3:**
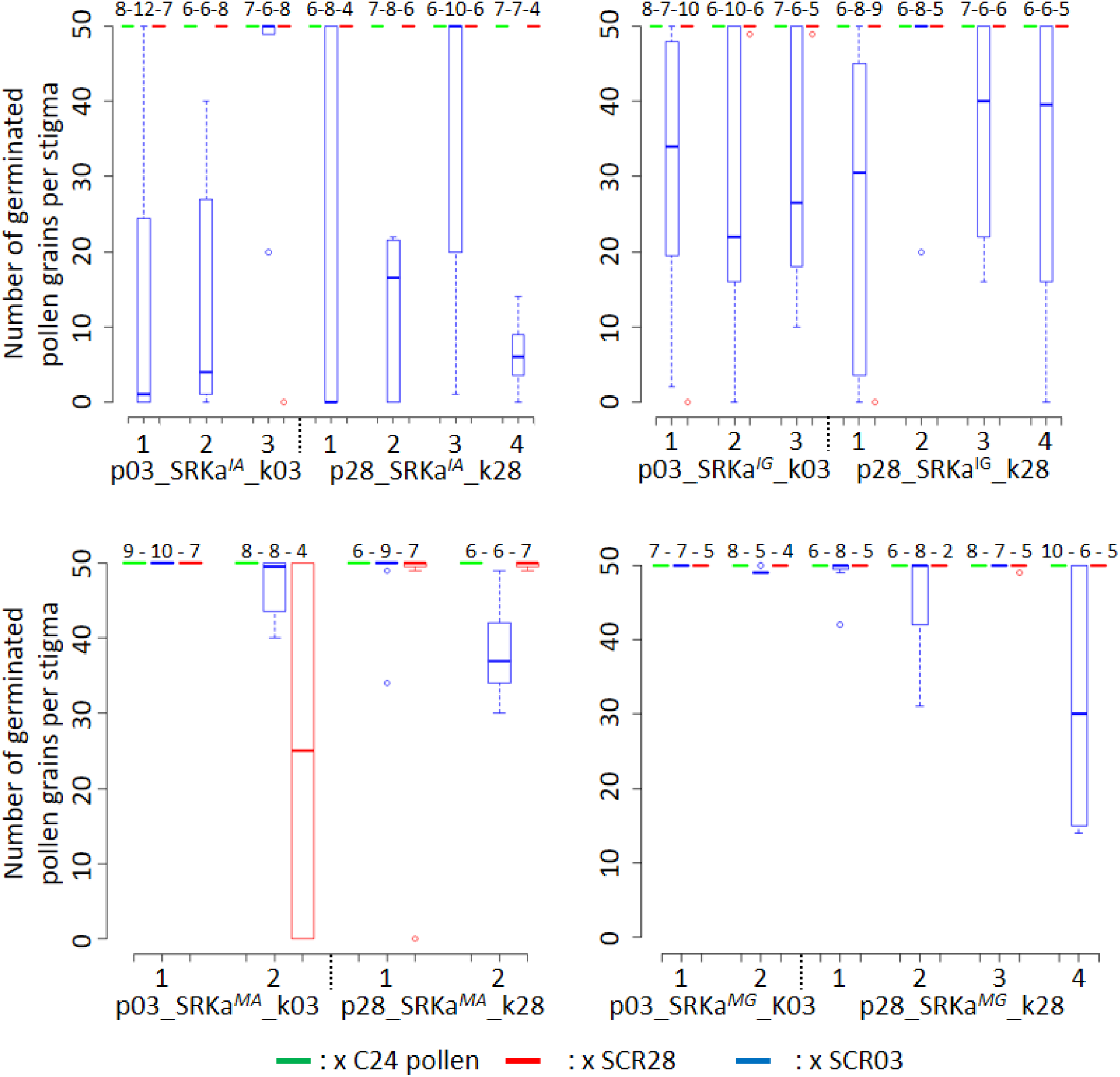
Incompatibility response of SRKa lines. Each SRKa line was pollinated with A. thaliana C24 (green), A. halleri S03 (blue) and S28 (red) pollen. The number of replicate pollinated pistils is indicated above each boxplot. The horizontal bar represents the median, the box delimitates the 25 and 75% percentiles, the bottom and top whiskers both represent 25% extreme percentiles and outliers are represented by individual dots.

With a single exception (line *p03_SRKa^MA^_k03_2)*, S28 pollen induced no consistent SI rejection in any of the lines expressing the different versions of the ancestral receptor, and was indistinguishable from the compatible control (C24, Fig 3). In stark contrast, SCR03 pollen induced robust incompatibility reactions in the vast majority (12 of 14) of the ancestral lines with amino-acid I at position 208 (*SRKa^IA^* and *SRKa^IG^*, Fig 3). Hence, we conclude that the S03 haplotype has retained the ancestral recognition specificity largely unaltered, while the S28 haplotype underwent substantial modification and represents a novel recognition specificity, leading to functional diversification.

Reactivity of ancestral lines was largely independent of the sequence context into which we introduced the different versions of the ancestral S-domain (Fig S3). Similarly, the identity of amino acid 305 of the reconstructed sequence had no impact on the response, as lines with either A or G produced rejection reactions in equal proportions. In contrast, only four of the lines with amino acid M at position 208 (*SRKa^MA^* and *SRKa^MG^*) displayed incompatibility reactions, and these reactions were weak (Fig 3). Hence, substitution of a single amino acid at position 208 (I to M), seems to be sufficient to almost fully inactivate receptivity of the ancestral receptor towards SCR03.

### Structural determinants of SRK specificity

We next sought to identify the structural determinants of the S03 and S28 specificities. We used the structural model of the interaction complex formed by SCR and SRK (Ma et al., 2016) to compare cognate and non cognate complexes in terms of number and nature of atomic contacts and predict how the ancestral receptor is expected to interact with either SCR03 or SCR28. As previously demonstrated by Ma et al. (2016), SCR binding induces homodimerization of the extracellular domain of SRK (eSRK) forming eSRK:SCR heterotetramers composed of two SCR molecules bound to two SRK molecules (Ma et al., 2016). The switch in recognition specificity occurred along the branch leading to *SRK28*, and therefore must involve some of the 29 amino acid differences along this branch. In contrast, the 31 amino acid differences that accumulated along the branch leading to *SRK03* have not fully altered the recognition specificity. Overall, we find a slightly higher number of atomic contacts between the SCR and SRK molecules in the SRK03-SCR03 cognate complex (1,543 contacts over 91 and 62 distinct SRK and SCR amino acids respectively) than in the SRK28-SCR28 cognate complex (1,395 contacts over 89 and 57 distinct SRK and SCR amino acids respectively, Table S1). As already noted by Ma et al. (2016) in Brassica and in line with the distribution of positively selected sites along the sequence (Castric and Vekemans 2007), in both cases the interaction interface was mostly concentrated around the three “hypervariable” portions of the SRK protein (Fig 4).

**Figure 4:**
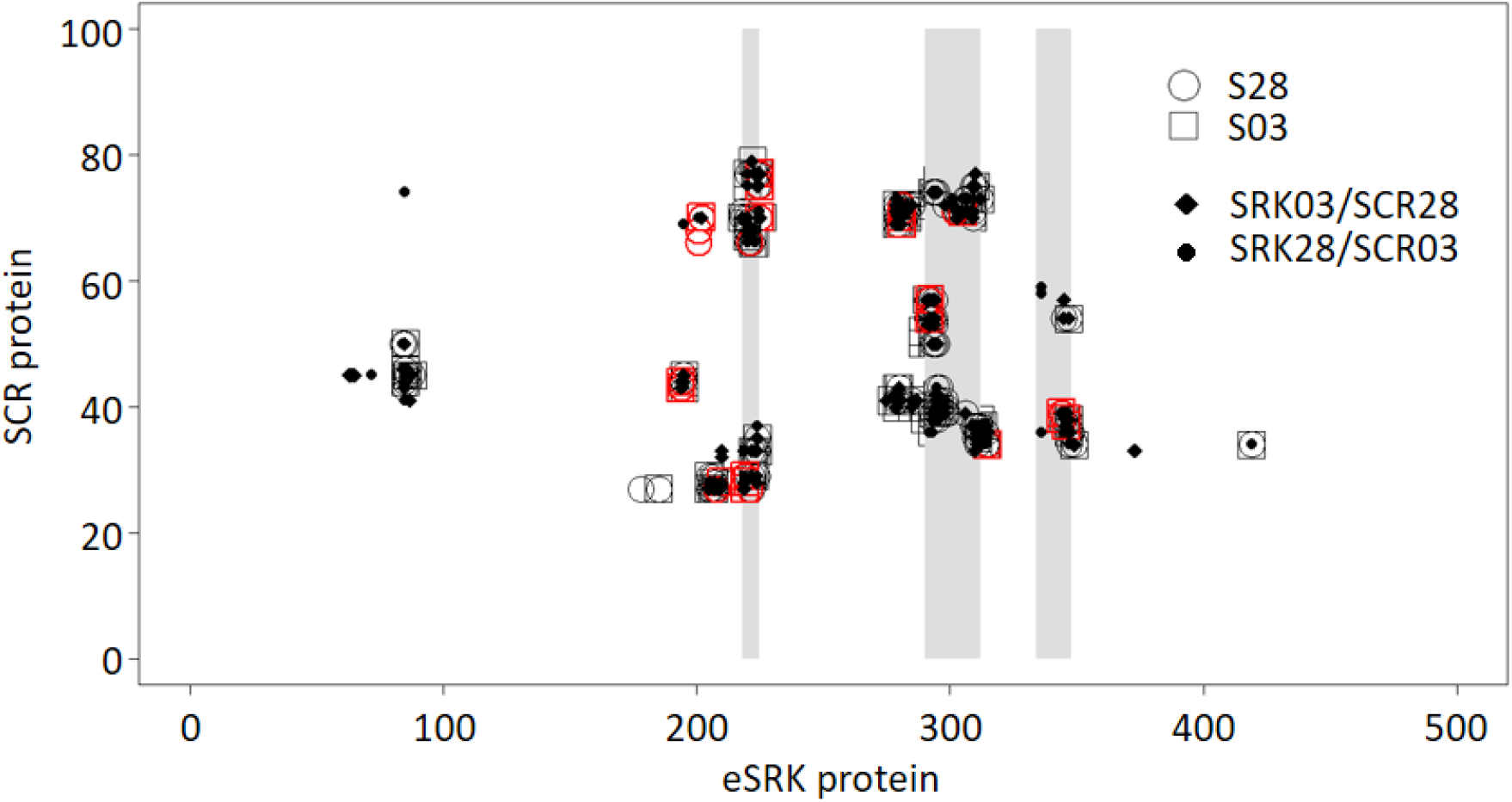
Map of atomic contacts from structural modelling of S03 and S28 complexes. The X and Y axis represent the eSRK and SCR protein respectively. Open circles and squares represent amino-acids contacts between eSRK and SCR proteins in cognate S03 and S28 complexes respectively. Red symbols represent SRK amino-acids that differ between SRK03 and SRK28. Full diamonds and dots correspond to amino-acids contacts in non-cognate SRK03/SCR28 and SRK28/SCR03 complexes respectively. Hypervariable regions 1, 2 and 3 of eSRK are represented by vertical grey bars at position 219-225, 290-312 and 334-348 respectively (Ma et al., 2016).

Contrary to our expectation however, the non-cognate complexes did not differ substantially from the cognate complexes on the basis of their total number of atomic contacts (Fig 4). Specifically, the SRK03/SCR28 and SRK28/SCR03 non-cognate complexes were predicted to establish a total of 1,546 and 1,328 contacts, respectively, between atoms of their SCR and SRK molecules, very close to the 1,543 and 1,395 predicted contacts for the SRK03/SCR03 and SRK28/SCR28 cognate complexes (Table S1). They were also not characterized by any obvious steric clashes that would prevent the proper docking of SCR and SRK. Hence, the overall stability of the complex does not seem to be the primary determinant of recognition specificity. Consequently, the specificity of the interaction must be a function of some qualitative features rather than of the quantitative strength of the interaction.

While the majority of amino-acids retained unchanged atomic contacts in cognate vs. non-cognate interactions, using a 5% threshold we identified six amino-acids positions of the eSRK protein that displayed important differences in terms of contacts they establish with SCR in the cognate vs. the non-cognate complexes (Fig S8B, S8C, S8E and S8F). Four of these amino-acids positions also displayed very contrasted interactions in the S03 vs. the S28 complexes (residues S201, R220, R279, W296, Fig S8A and S8D). For instance, the R residue at position 279 of SRK establishes three times more contacts with SCR in the SRK03/SCR03 than in the SRK28/SCR28 complex (Fig 5, Fig S8A). These four residues are therefore prime candidates for the determination of binding specificity. Intriguingly, three of the six amino-acids were identical between SRK03 and SRK28, and yet interacted in sharply different ways with their respective SCR ligands. For instance, while identical between the SRK03 and SRK28 sequences, residues W296 and R279 established three to four times more contacts in the S03 than in the S28 complex, and the R220 residue is involved in twice as many contacts in the S28 than in the S03 complex (Fig 5, Fig S8A). This suggests that involvement of these residues in the activity of the complex is mediated by substitutions at other positions along the protein sequence, for instance by displacing them spatially (Fig 5).

**Figure 5:**
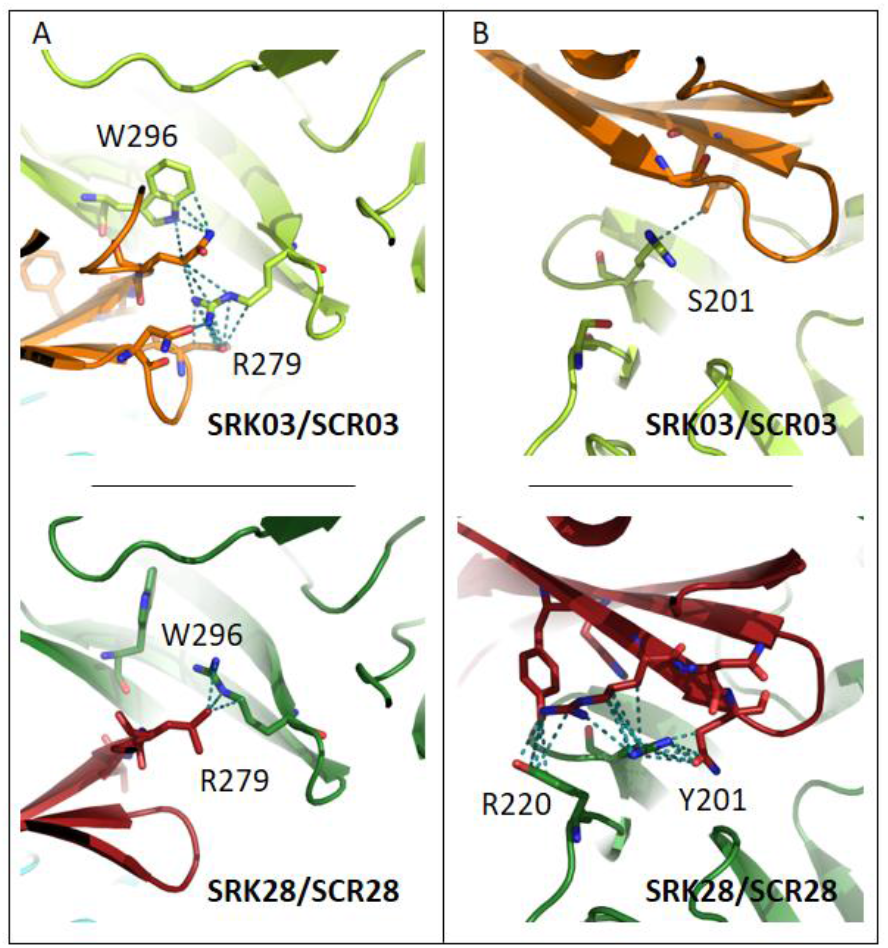
Four amino-acid residues establish contrasted patterns of atomic contacts between SCR and SRK in the two cognate complexes. In **(A)**, residues R279 and W296 of eSRK establish a large number of atomic contacts with SCR in the S03 complex (upper panel), but a very low number of contacts in the S28 complex (lower panel). The situation is reversed in **(B)**, where residues S201 and R220 of eSRK establish a large number of atomic contacts with SCR in the S28 complex (lower panel), but a very low number of contacts in the S03 complex (upper panel). SRK chains are coloured in light and dark green for SRK03 and SRK28, respectively; SCR chains are coloured in orange and red for SCR03 and SCR28, respectively. Amino acid residues are shown in stick representation, with dotted lines indicating atom pair contacts below 4 Å, excluding hydrogen atoms. Note that for clarity a more stringent threshold was used to define atomic contacts here (4 Å) than in Fig S8 and S9 (where a 5 Å threshold was used for a more comprehensive analysis), but the results are qualitatively similar.

We then compared the predicted binding features of the functional form of the ancestral receptor (SRKa^*IA*^) with both SRK03 and SRK28. When complexed with SCR03, 46 SRKa^IA^ interacting pairs of amino acids were identical with SRK03, whereas only 24 were identical with SRK28. Similarly, when complexed with SCR28, 37 SRKa^*IA*^ interacting pairs of amino acid residues were identical with SRK03, but only 18 were identical with SRK28 (Fig 6). This trend is also visible at the level of individual atomic contacts and at the level of involved amino acids both in SRK and SCR molecules (Fig S9). Hence, even though there is no quantitative difference in how strongly SRK03 and SRK28 bind SCR03 and SCR28 (Table S1), the amino acids that come into contact in the two complexes are not identical, and the contacts established by SRKa^*IA*^ with SCR qualitatively resemble more those established by SRK03 than those established by SRK28.

**Figure 6:**
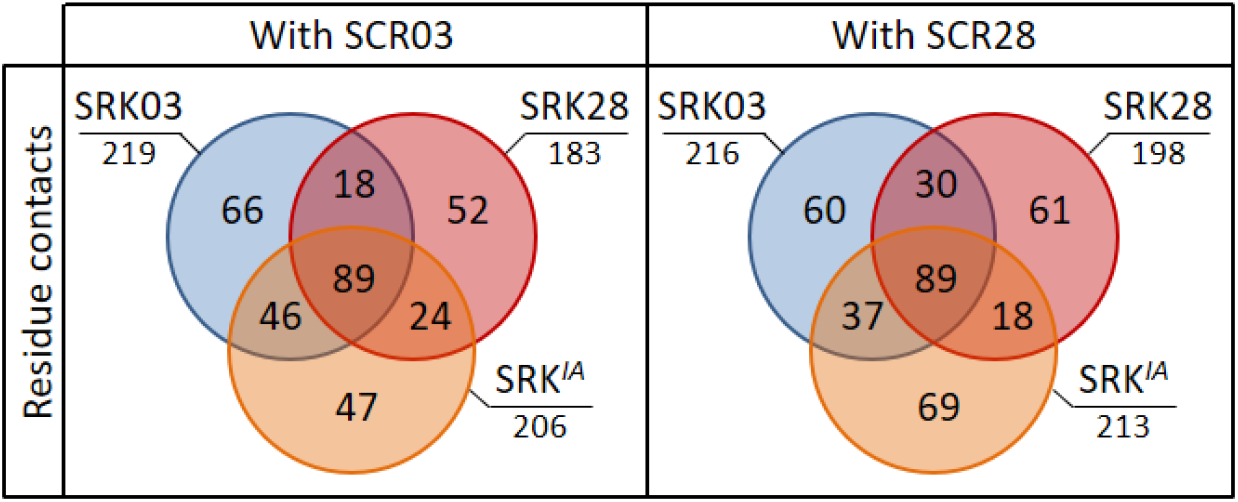
Comparison of predicted binding properties of SRK03, SRK28 and SRKa^IA^ receptors when forming a complex with either the SCR03 or the SCR28 ligand. Residues in contact between SRKa^IA^ and SCR are more often also in contact between SRK03 and SCR than between SRK28 and SCR. The Venn diagram indicates the number of contacts between amino acid residues that are shared or specific across variants of the SRK receptor and SCR.

Overall, asymmetrical diversification of the ancestor into the phenotypically distinct S03 and S28 specificities is therefore associated with structural changes of the SRK receptor and the way it interacts with its SCR ligand. Despite similar levels of overall sequence divergence between SRKa and either SRK03 or SRK28, the binding properties of SRK03 have remained similar to those of SRKa^*IA*^, while in contrast the way SRK28 is predicted to interact with the SCR ligand has diverged from its ancestral state, consistent with the phenotypic switch that has occurred along this lineage.

## Discussion

Receptor-ligand interactions are widespread and biologically important, yet the processes by which they become diversified in the course of evolution have remained poorly understood. Here, using an ancestral resurrection approach, we showed that the receptor controlling the female SI recognition specificity in *A. halleri* evolved asymmetrically by accumulation of specificity-altering mutations along a single lineage, leaving the other descendent lineage functionally unchanged. We identify structural modifications of the receptor-ligand complex that are associated with this functional divergence.

### Ancestral resurrection in a higher organism

Ancestral resurrection approaches have been successfully used to decipher evolutionary processes in a number of biological systems including enzymes (Starr, Picton, and Thornton 2017; Tufts et al., 2015), visual pigments (Chang et al., 2002) or transcription factors (Pougach et al., 2014). However, an essential limitation in these studies is that the ancestral protein function is typically determined *in vitro*, strongly limiting biological realism of the reconstructed function. To our knowledge, a single study conducted a comparable *in vivo* ancestral resurrection approach in a higher organism, in the context of functional evolution of the alcohol dehydrogenase protein (ADH) in *Drosophila* lineages (Siddiq et al., 2017). Such whole organism studies are biologically more relevant, as they allow to focus on integrated phenotypes (here the self-incompatibility reaction, which has a strong and direct link with organismal fitness) rather than on highly reductionist phenotypes (such as in vitro assays of protein activity).

### Asymmetrical structural and functional divergence

Functional diversification of receptor-ligand interactions is a common process that takes place over long evolutionary times. However, in most cases we remain far from a predictive understanding of how the specific evolutionary forces imposed on a given interacting pair of proteins at the molecular level translate into a given process of functional diversification. As a result, little is known about how diversification happens in Nature, both in terms of whether some evolutionary pathways are preferentially followed over others and in terms of the molecular modifications involved. Here, by focusing on the SI system of *A. halleri*, we demonstrate conservation of recognition specificity between an ancestral SRK receptor and one of its two descendant lineages (S03), while pollen from the other descendant lineage (S28) triggered no substantial SI response when deposited on pistils expressing the ancestral specificity. Hence, S-allele diversification of the two S-alleles from their common ancestor proceeded through asymmetrical functional divergence, with one allele having largely retained its ancestral specificity and the other one having acquired a novel recognition specificity that was previously absent. In analogy with macroevolution models, the diversification of S-locus specificities thus seems to proceed through cladogenetic change rather than anagenetic change (e.g. Goldberg & Igic, 2012).

Overall, our results thus unambiguously reject two previously proposed models of SI specificity diversification. The model of gradual divergence of pairwise SRK-SCR affinities along lineages (Chookajorn et al., 2004) is rejected because both descendent alleles would have been functionally distinct from the ancestor. The model of promiscuous dual-specificity intermediates (Matton et al., 1999) is rejected because Uyenoyama & Newbigin (2000) have shown that a dual-specificity haplotype can be maintained in the population only in the absence of the ancestral specificity, while we show here that the ancestral specificity (S03) was indeed present. In contrast, the remaining scenario (in which diversification proceeds through transient self-compatible intermediates; Gervais et al., 2011; Uyenoyama et al., 2001) is compatible with our observation of long-term maintenance of functional specificities.

### Long-term maintenance of SI specificity rather than rapid turnover

Models of SI evolution suggested that turnover of recognition specificities may be common over evolutionary times (Gervais et al., 2011). The observation that the ancestral SRKa^*IA*^ recognition specificity was stably maintained along the SRK03 lineage, demonstrates that no turnover event occurred over a substantial period of time (i.e. since S03 and S28 started to diverge from one another, *ca*. a few million years ago). The long-term maintenance of S-haplotype specificities was previously hinted by the comparison of chosen *A. halleri* and *A. lyrata* S-lineages with their *A. thaliana* orthologs. Specifically, some *A. thaliana* accessions with haplogroup A were able to reject *A. halleri* pollen expressing the cognate AhSCR04 specificity (Tsuchimatsu et al., 2010), despite about 6 million years of divergence (Hohmann et al., 2015). Similarly, an overexpressed version of *A. thaliana SCR* haplogroup C was also able to elicit a SI reaction on pistils expressing the cognate *A. lyrata* AlSRK36 specificity (Dwyer et al., 2013). However, beside the fact that these were trans-specific comparisons, *A. thaliana* has become a predominant selfer due to alterations of SI components (Bechsgaard et al., 2006; Shimizu and Tsuchimatsu 2015), so our results provide the first direct test of the hypothesis that recognition specificities remain stable rather that undergo rapid and recurrent turnover.

Despite the obvious conservation of recognition specificity between SRKa^*IA*^ and SRK03, we note that the SI response towards the reconstructed SRKa was overall weaker and more variable across replicate lines than towards the native SRK constructs. A first possibility is that this partial response is due to uncertainty in the ancestral receptor reconstruction. Ancestral reconstruction is based on probabilistic inference, and our results show that a single amino acid modification (I vs M at position 208) can entirely compromise the SI response. It is therefore possible that the resurrected ancestor had a partly diminished capacity to elicit the SI reaction because of imperfect reconstruction. A second possibility however, is that this partial response is due to parallel changes to the SCR ligand, suggesting that amino acid substitutions occurring between SRKa and SRK03 have fine-tuned the SRK03/SCR03 interaction. Resurrection of the ancestral SCRa ligand will now be needed to solve this issue.

### Structural determinants of the shift in binding specificity of a receptor-ligand interaction

Structural modelling of the cognate vs non-cognate complexes suggested that specificity of the receptor/ligand interaction depends on qualitative features rather than quantitative strength of the interaction. In other words, a ligand is predicted to bind its non-cognate just as well as its cognate receptor, but it only activates the proper one by interacting with a different set of amino acid residues at the binding interface. The importance of such allosteric activation has been discussed in the context of protein-protein interactions (Kang et al., 2015) as well as of the binding of transcription factors to their DNA binding sites (Gronemeyer and Bourguet 2009) and may be a general feature of molecular interfaces (Ma et al., 2010). In our system, the model of allosteric activation points to a limited number of amino-acid residues encoding recognition specificity and will now need to be tested by direct measurements of binding affinity, in line with Ma et al. (2016). Specifically, it will be interesting to determine experimentally whether the allosteric model proposed here also applies to more divergent pairs of SCR-SRK variants, which are more typical of the SI system given the long-term maintenance of this balanced polymorphism.

## Materials and Methods

### Phylogeny-based ancestral SRK resurrection

#### Sequence collection

In order to reconstruct the ancestral sequence of the *AhSRK28* and *AhSRK03* SRK alleles, we first collected 17 full or partial sequences of the *SRK* first exon belonging to the haplogroup B/2. Seven allele sequences belonged to *A. halleri* (*AhSRK03, AhSRK08, AhSRK09, AhSRK19, AhSRK23, AhSRK27, AhSRK28*), six to *A. lyrata* (*AlSRK06, AlSRK08, AlSRK14, AlSRK18, AlSRK29, AlSRK39*) and four to *Capsella grandiflora* (*CgrSRK1, CgrSRK4, CgrSRK5, CgrSRK6*). For the *AhSRK19, AhSRK23 and AhSRK27* alleles the *SRK* exon1 partial sequence was supplemented by Sanger sequencing. We added three sequences from the *SRK* haplogroup A3/3 as outgroups (*AhSRK04, AhSRK10, AhSRK29*). Accession numbers for the sequences are reported in Table S2. After preliminary analyses and in order to obtain a well-supported phylogenetic tree we removed three fast evolving sequences: *CgrSRK6, AhSRK08* and *AlSRK29*.

#### Phylogenetic analysis

Sequences were aligned with MACSE v1.2 (Ranwez et al., 2011). Based on the 1338 base pairs alignment, SRK phylogenetic trees were built with both Maximum Likelihood (ML) (PHYML 3.0, Guindon et al., 2010) and Bayesian methods (MrBayes 3.2.4, Ronquist & Huelsenbeck, 2003). For the ML analysis the model used was chosen according to jModelTest 2.1.10 (Darriba et al., 2012) using the Bayesian Information Criterion (BIC) (TPM3uf+Γ). Node stability was estimated by 100 non-parametric bootstrap replicates (Felsenstein 1985). For the Bayesian inference (BI), a codon model with a GTR+Γ+I model of substitution was used; two runs of four Markov chains were calculated simultaneously for 400,000 generations with initially equal probabilities for all trees and a random starting tree. Trees were sampled every 10 generations, and the consensus tree with posterior probabilities was calculated after removal of a 25% burn-in period. The average standard deviation of split frequencies between the two independent runs was lower than 0.01. The resulting topology was used to infer *SRKa* (Fig S10).

#### Ancestral SRK reconstruction

The ancestral reconstruction was conducted with the codeml program of the PAML 4.8 package (Yang 2007) under six different models: M0, M0-FMutSel0, M0-FMutSel, M3, M3-FMutSel0, M3-FMutSel. M0 refers to a one-ratio model, which assumes a single ω (d_n_/d_s_) across branches and sites (Goldman and Yang 1994), whereas M3 allows ω to vary across sites according to a discrete distribution (with 2 or 3 categories depending on the models) (Yang et al., 2000). The mutation-selection models incorporate parameters for mutation bias, with (FMutSel) or without (FMutSel0) selection on synonymous rate (Yang and Nielsen 2008). Models were compared using Likelihood Ratio Tests (LRT) and Akaike Information Criteria (AIC). The best fitting-model was found to be M3FMutSel (Table S3), therefore the ancestral amino acid sequence reconstructed with this model was used to generate transgenic lines (SRK^IA^). The ancestral amino acids inferred with the three “M3” models were identical except for three sites with relatively lower probability: 33 (A or V), 208 (I or M) and 305 (A or G) (Fig S5). Two of these sites were close (208) or within (305) hyper variable regions and therefore potentially functionally important. To take into account these uncertainties, three additional transgenic lines were generated with a different SRKa and tested: SRK^IG^, SRK^MA^ and SRK^MG^, in order to evaluate all four combinations of two amino acids at these two sites.

### Generation and selection of A. thaliana transgenic lines

We generated 12 series of *A. thaliana* C24 transgenic plants (*AhSRK03; AhSRK28; AhSRK03p:GFP; AhSRK28p:GFP; p03_SRKa^IA^_k03; p28_SRKa^IA^_k28; p03_SRKa^IG^_k03; p28_SRKa^IG^_k28; p03_SRKa^MA^_k03; p28_SRKa^MA^_k28; p03_SRKa^MG^_k03* and *p28_SRKa^MG^_28*). All DNA amplifications were performed with the primeSTAR^®^ DNA polymerase (Takara, Japan) and all constructs were validated by SANGER sequencing. We used gateway vectors^®^ (Life Technologies, USA) for expression of transgenes in *A. thaliana. AhSRK03, AhSRK28, proAhSRK03* and pro*AhSRK28* were amplified using attB1 and attB2 containing primers (Table S4) from BAC sequences of *A. halleri* S03 and S28 (KJ772378-KJ772385 and KJ461475-KJ461478 respectively, Goubet et al., 2012). Amplification products were inserted by BP recombination into the pDONR 221 plasmid. Promoters pro*AhSRK03* and pro*AhSRK28* designed for p03_SRKa_k03 and p28_SRKa_k28 constructs respectively were amplified with attB4 and attB1r containing primers (Table S4). Amplification products were inserted by BP recombination into a pENTR-P4-P1R plasmid. Kinase domains of AhSRK03 and AhSRK28 were amplified with attB2r and AttB3 containing primers (Table S4). Amplification products were inserted by BP recombination into a pENTR-P2R-P3 plasmid. Constructs expressing native SRK (*AhSRK03* and *AhSRK28*) and GFP expressing constructs (*AhSRK03p:GFP* and *AhSRK28p:GFP)* were generated by LR reaction between entry clones and the destination vectors pK7WG or pKGWFS7.0. SRKa constructs were generated by LR triple recombination into the pK7m34GW destination vector with i) attL4 and attR1 promoters entry clones, ii) synthesized ancestral S-domain surrounded by attL1 and attL2 sequences and iii) attR2 and attL3 kinase domain entry clones. Details of the molecular constructs are displayed in Fig S11. *A. thaliana* plants were grown under 16h/20°C day, 8h/18°C night and 70 % humidity greenhouse conditions. When plants displayed approximately 20 flowers, transformation was performed by *Agrobacterium tumefaciens*-mediated floral deep according to Logemann et al., 2006). After seeds harvesting, single insertion homozygous lines were selected *via* multiple rounds of antibiotic selection on selective medium as described in Zhang et al., 2006).

### Microscopy

GFP expression in *AhSRK03p:GFP* and *AhSRK28p:GFP* and the results of cross pollination experiment were monitored under UV light using an Axio imager 2 microscope (Zeiss, Oberkochen, Germany) coupled with a HXP120 Light source (LEJ, Jena, Germany). Pictures were taken with an Axiocam 506 color camera (Zeiss, Oberkochen, Germany) and read with the Zeiss ZEN 2 core software package. For *GFP* expression, floral buds, pieces of leaves and floral hamp cross sections from *AhSRK03p:GFP* and *AhSRK28p:GFP* were observed. For pollination assays, the number of germinated pollen grains per stigma was counted after aniline blue staining up to 50 pollen tubes.

### Aniline blue staining

To evaluate the SI response of transgenic lines, floral buds of developmental stage 9-11 were emasculated at day-1. At day-2, they were pollinated with frozen *A. halleri* pollen expressing S03 or S28 specificity. Pistils were harvested 6h after pollination and fixed in FAA (4 % formaldehyde, 4 % acetic acid, ethanol) overnight. On day-4, pollinated stigmas were washed three times in water, incubated 30 min in NaOH 4M at room temperature, washed three times in water and conserved for 30 min in the last bath. Finally, pistils were incubated overnight in aniline blue (K3PO4 0.15 M, 0.1 *%* aniline blue) and mounted between microscopic slides at day-5.

### Expression analysis

RNA extraction was performed onto stages 11-14 floral buds with the nucleospin RNA Plus Extraction kit (Macherey Nage, GmbH & Co. DE). For each line, floral buds were harvested onto three different individuals. cDNA synthesis was performed using RevertAid Reverse Transcriptase (Thermo Fisher Scientific, Massachusetts, USA). 1μg of extracted RNA was mixed with 0.2μg of Random hexamer primer, psq H2O 12.5μL. After incubation at 65°C, 4μL of 5X reaction buffer; 0,5μL of Riboblock RNase inhibitor (Thermo Fisher Scientific, Massachusetts, USA), 10 mM of each dNTP and 200 U of RevertAid Reverse Transcriptase were added and incubated firstly at 25 °C and then 10 min at 42 °C. cDNA quantification was performed in triplicate using LightCycler 480 Instrument II (Roche molecular systems, Inc, Pleasanton, USA). cDNA amplification of *SRK* fragments was performed with forward primer located on the S-domain (For: AGGAATGTGAGGAGAGGTGC) and the reverse primer located on the second exon (Rev: TCCTACTGTTGTTGTTGCCC). According to Liu et al. (2007), Ubiquitin was used as housekeeping gene (For: CTGAGCCGGACAGTCCTCTTAACTG; Rev: CGGCGAGGCGTGTATACATTTGTG).

### Interacting protein modelling

All structural models were created using MODELLER (Fiser, Do, and Šali 2000; Šali and Blundell 1993), applying a template-based docking approach with the eSRK9:SCR9 X-Ray structure (Ma et al., 2016) as reference. Input sequence alignments were created using multiple sequence alignments of all relevant SRK sequences and of all relevant SCR sequences. The resulting models were subsequently optimized using the GalaxyRefineComplex web service (Heo, Lee, and Seok 2016).

The top models for each receptor-ligand pair were then considered and the number of atomic contacts between chains was counted using a 5 Å atom-atom cut-off. Hydrogen atoms were excluded from the calculation. Visualization was done, and for clarity images were made with a 4 Å atom-atom cut-off using the PyMol molecular graphics system, version 1.7.2.1, Schrodinger LLC.

## Acknowledgements

This research received support through a grant from the European Research Council (NOVEL project, grant #648321). The authors thank the French Ministère de l’Enseignement Supérieur et de la Recherche, the Hauts de France Region and the European Funds for Regional Economical Development for their financial support to this project.

## Supplementary Materials

**Figure S1:**
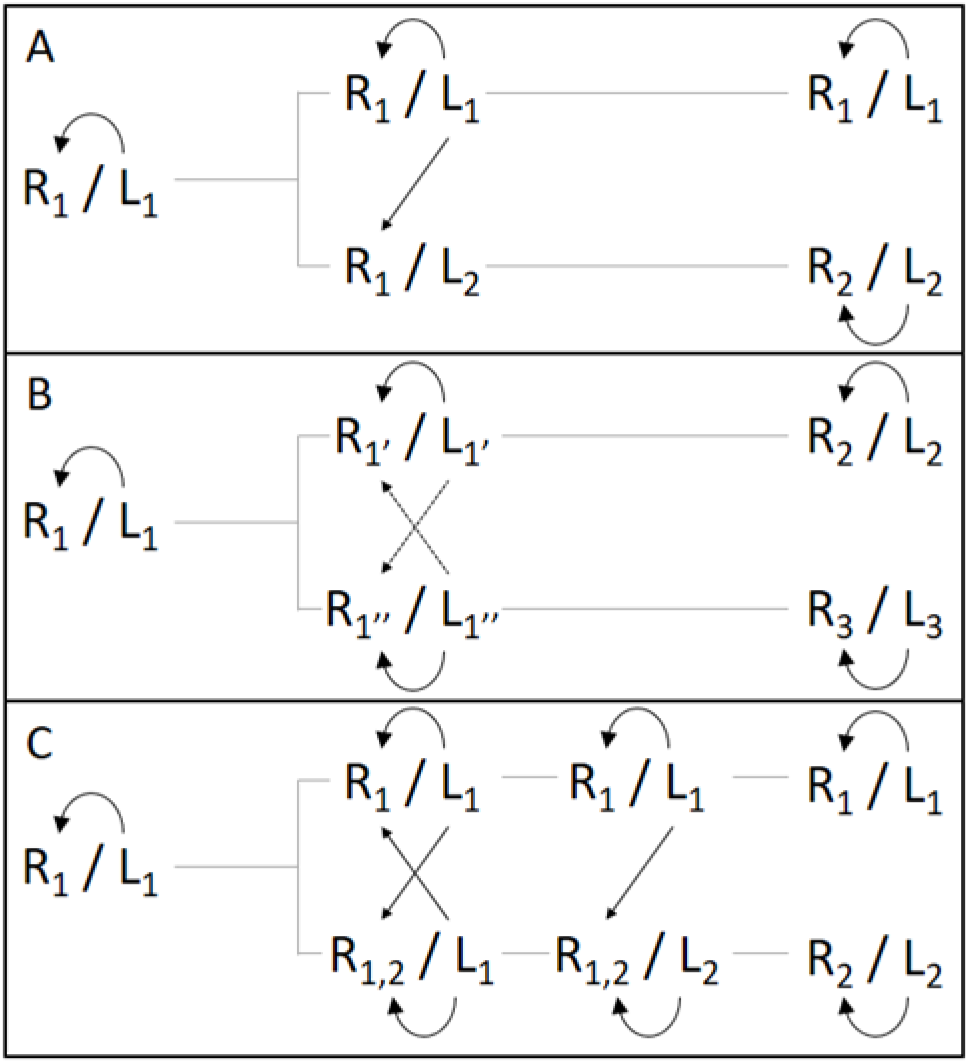
Three models for the emergence of new self-incompatibility specificities. Docking interaction between ligands and receptors are represented by oriented arrows. **A**. In the compensatory mutation model, a non-functional self-compatible intermediate (R_1_/L_2_) segregates transiently and is rapidly compensated, creating a novel specificity (R_2_/L_2_), while the ancestral one (R1/L1) remains unchanged over time. **B**. In the turnover model, slight functional variants with different affinity between them (R_1’_/L_1’_ and R_1”_/L_1”_) segregate in the population and give rise to new specificities (R_2_/L_2_ and R_3_/L_3_). Dotted lines correspond to weaker affinity interactions. **C**. In the promiscuous model, the intermediate receptor (R_1,2_) has widened up its specificity spectrum enabling it to recognize another potential ligand (L_2_) while maintaining its capacity to recognize its original ligand (L_1_). Emergence of the new ligand then favours narrowing of specificity the dual-receptor (R_1,2_ → R_2_)

**Figure S2:**
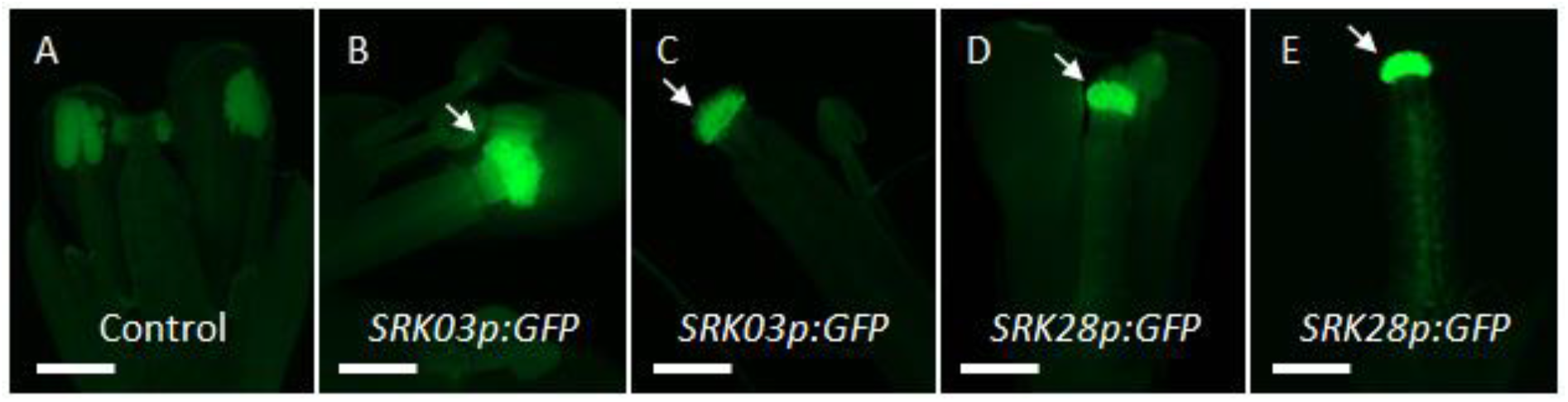
Localization of SRK expression by GFP florescence microscopy. **(A)** Flower of an untransformed plant (no SRK expression). **(B,C)** Flowers of two SRK03p:GFP transformed lines. **(C,D)** Flowers of two SRK28p:GFP transformed lines. Promoters of both SRK03 and SRK28 drive GFP expression specifically in stigmatic papillae cells (white arrows). Bar = 1mm.

**Figure S3:**
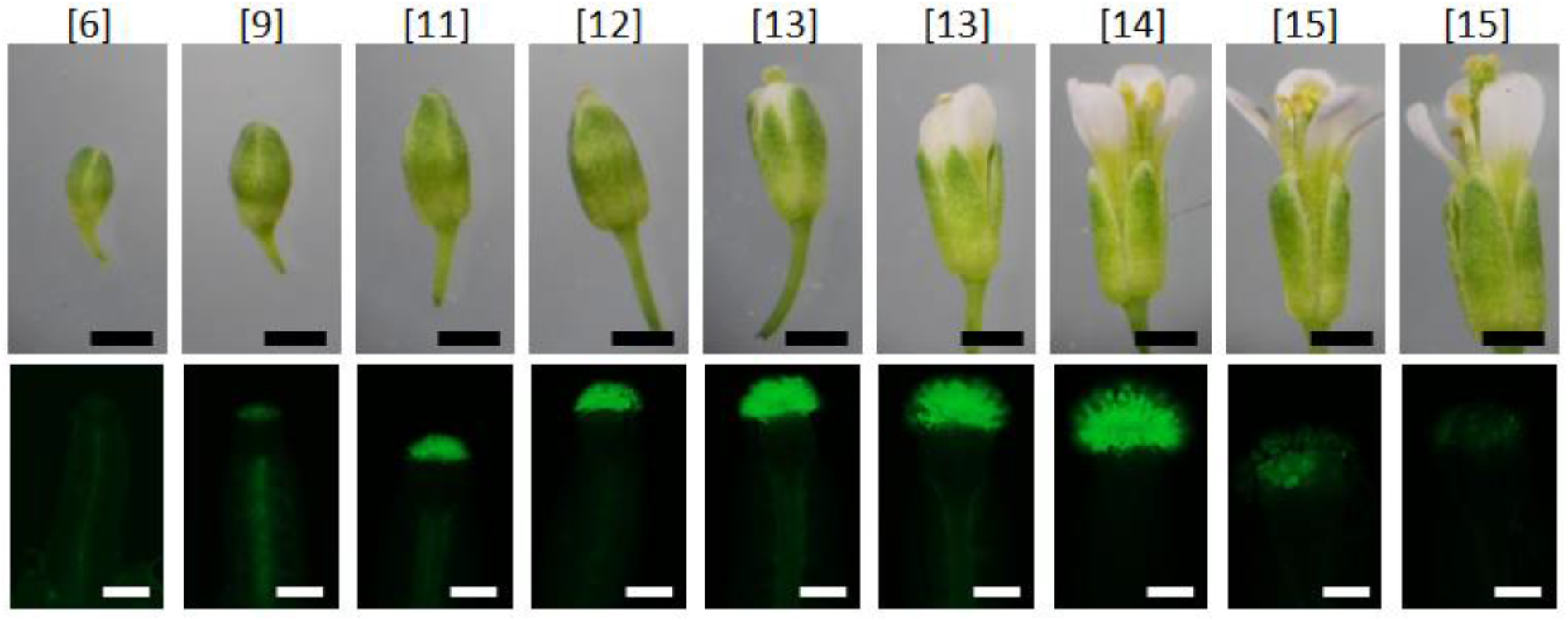
Pattern of proSRK03 activity along development stages of floral buds. Floral buds and pistils of SRK03p:GFP-transformed plants are observed under optical (above) and UV light (below), respectively. Developmental stages (Smyth et al., 1990) are indicated between brackets. SRK expression is located between stages 11 and 14, with a strong decrease of GFP fluorescence in mature flowers (old stage 15). Similar results were observed for the SRK28p:GFP lines (data not shown). No GFP signal was observed in any other part of the plant (inflorescence stems, leaves and non-reproductive floral organs, data not shown). Black bar = 1 mm, white bar = 0.5 mm.

**Figure S4:**
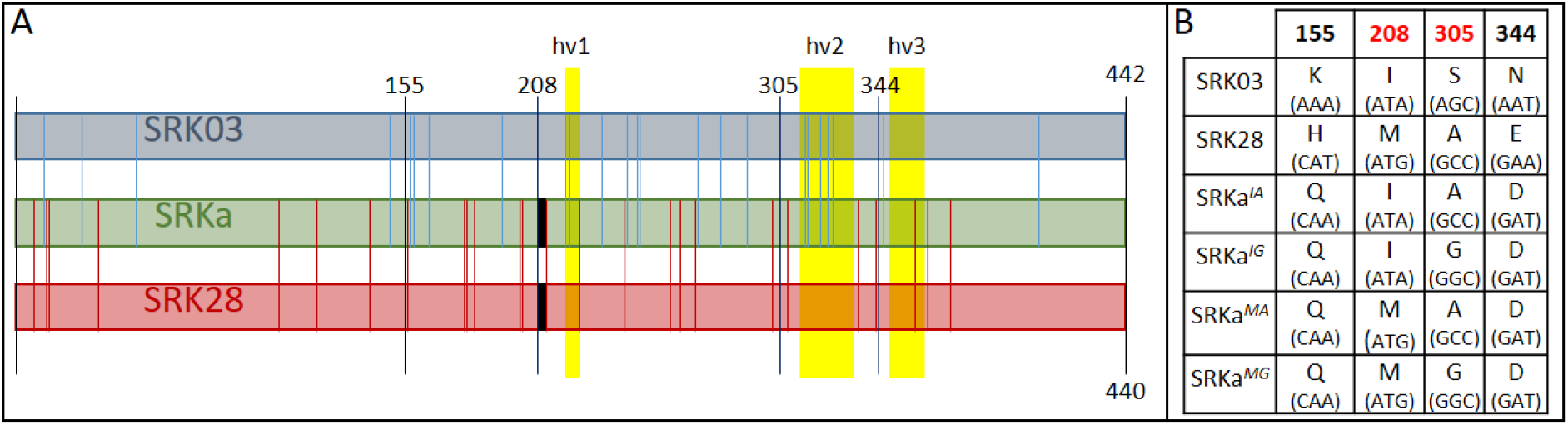
Comparison between the S-domain of AhSRK03 and AhSRK28 and their common ancestor. **(A)** Protein schematic representation of the S-domain sequence of AhSRK03, AhSRK28 and their common ancestor SRKa. Positions of identical amino acids between the three S-domains are not represented, positions of SRKa amino acids specific to SRK03 and SRK28 are represented by a vertical blue and red line respectively. Positions of hypervariable regions are represented by yellow rectangles. The position of the 2 amino acid gap presents in both AhSRKa and AhSRK03 is represented by black rectangles. Positions 208 and 305 for which the reconstruction was uncertain and positiosn 155 and 344 for which the aa identity is different in the two other receptors are represented in black. **(B)** Amino acid identity and codon sequence (in parentheses) for these four positions in the different SRK variants.

**Figure S5:**
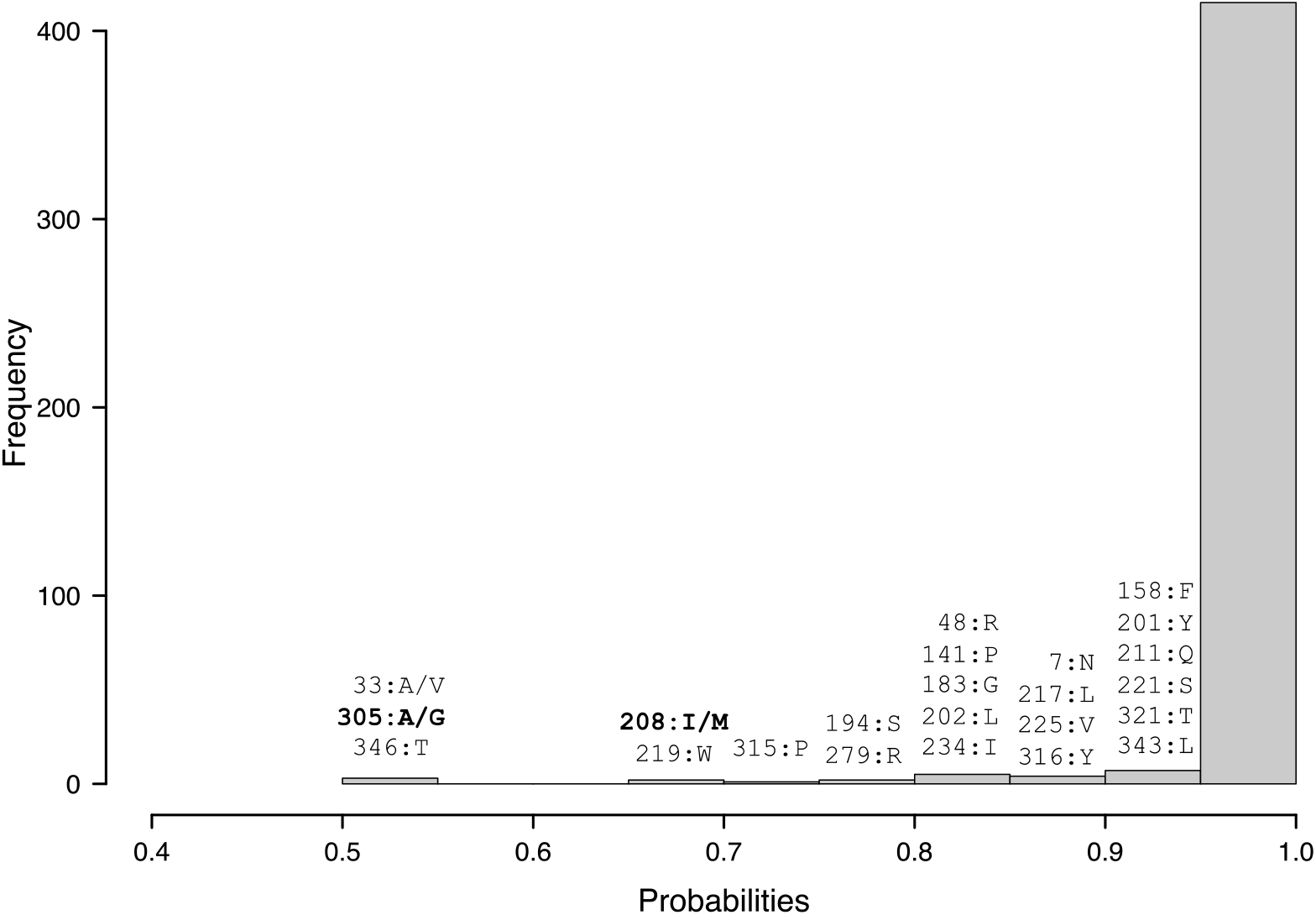
Histogram presenting the distribution of amino acid (aa) probability values obtained with the best fitting-model M3FMutSel. For each range of values smaller than 0.95, the nature and the position of the aa in the sequence are given. At three sites (33, 208, 305) the models used for the inferences led to different results. The variable sites between the 4 ancestral sequences used for *A. thaliana* transformation are in bold.

**Figure S6:**
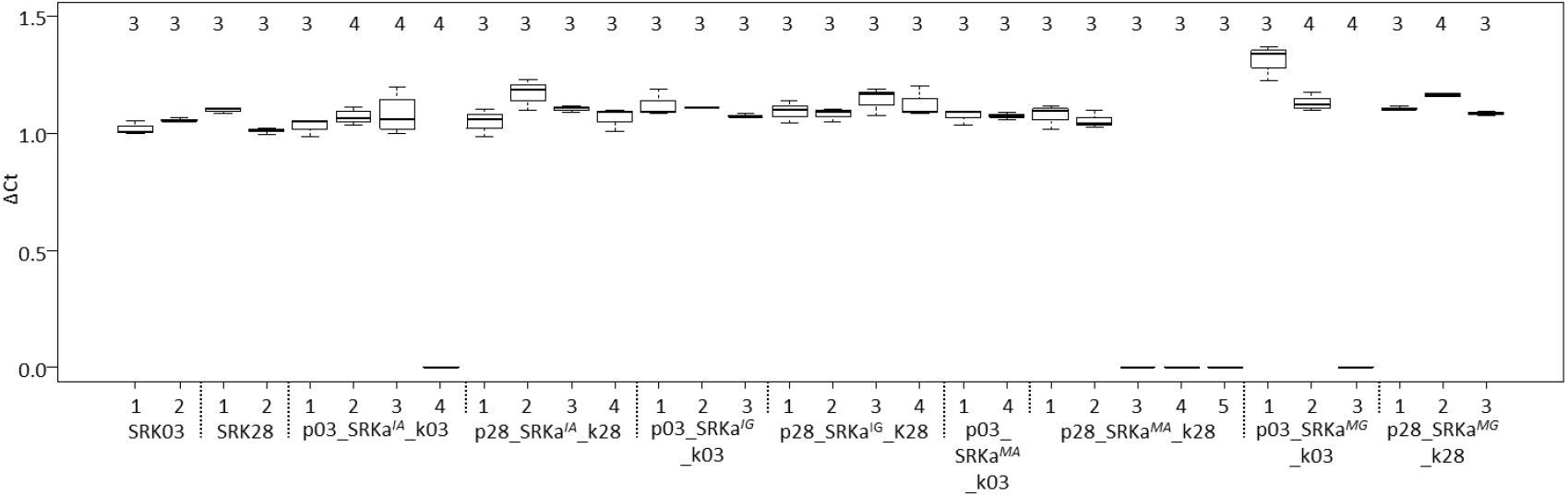
Transgene expression analysis of the different SRK-transformed lines. ΔCt (Cycle threshold) was calculated as the ratio of Ct-SRK on Ct-Ubiquitin. The number of biological replicates is indicated above each boxplot. For each biological replicate, measurements were realized in technical triplicates.

**Figure S7:**
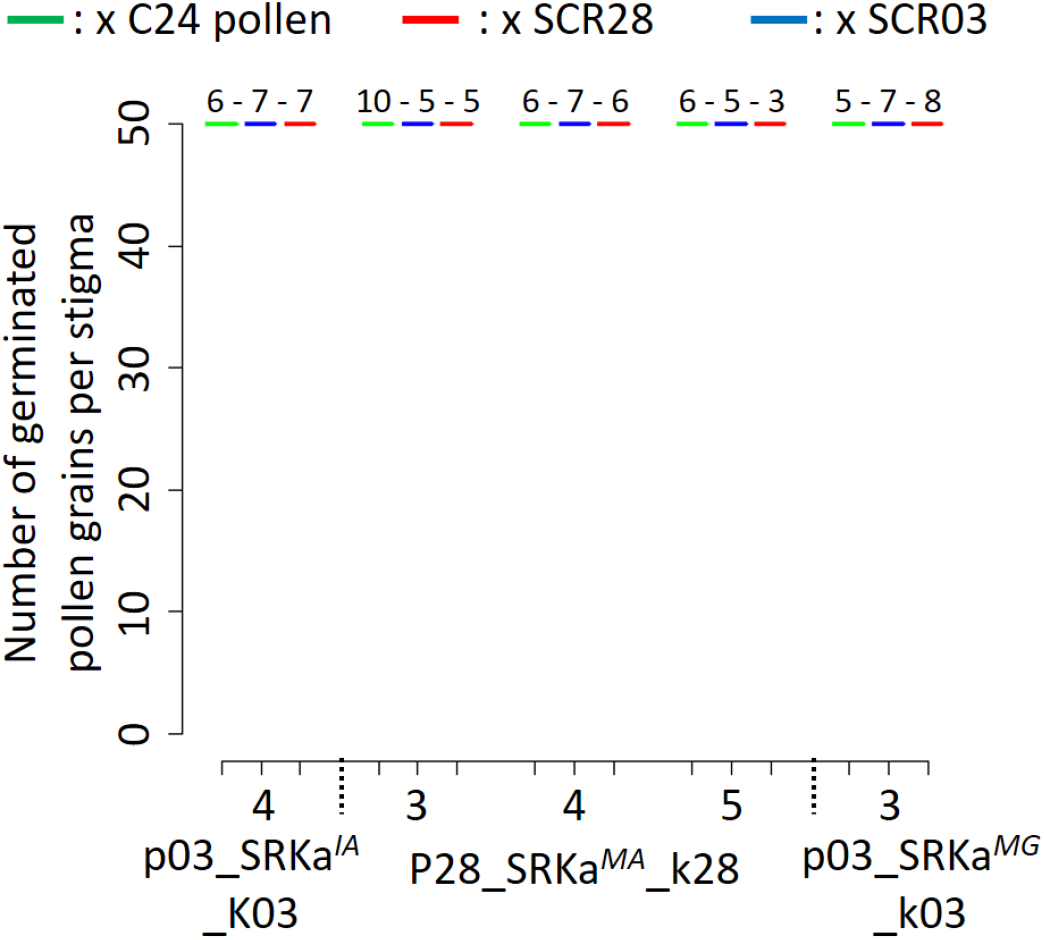
Quantification of incompatibility response of *A. thaliana* lines with SRKa transcript levels below detection threshold. Each SRKa line in S03 and S28 context were pollinated with *A. thaliana* C24 (green), A. halleri S03 (blue) and S28 (red) pollen. The number of pollinated pistils is indicated above each boxplot. These five lines with an undetectable level of transcripts appear compatible with the three pollen types, which is in agreement with the fact that they don’t express any SI receptor at their membrane surface.

**Figure S8:**
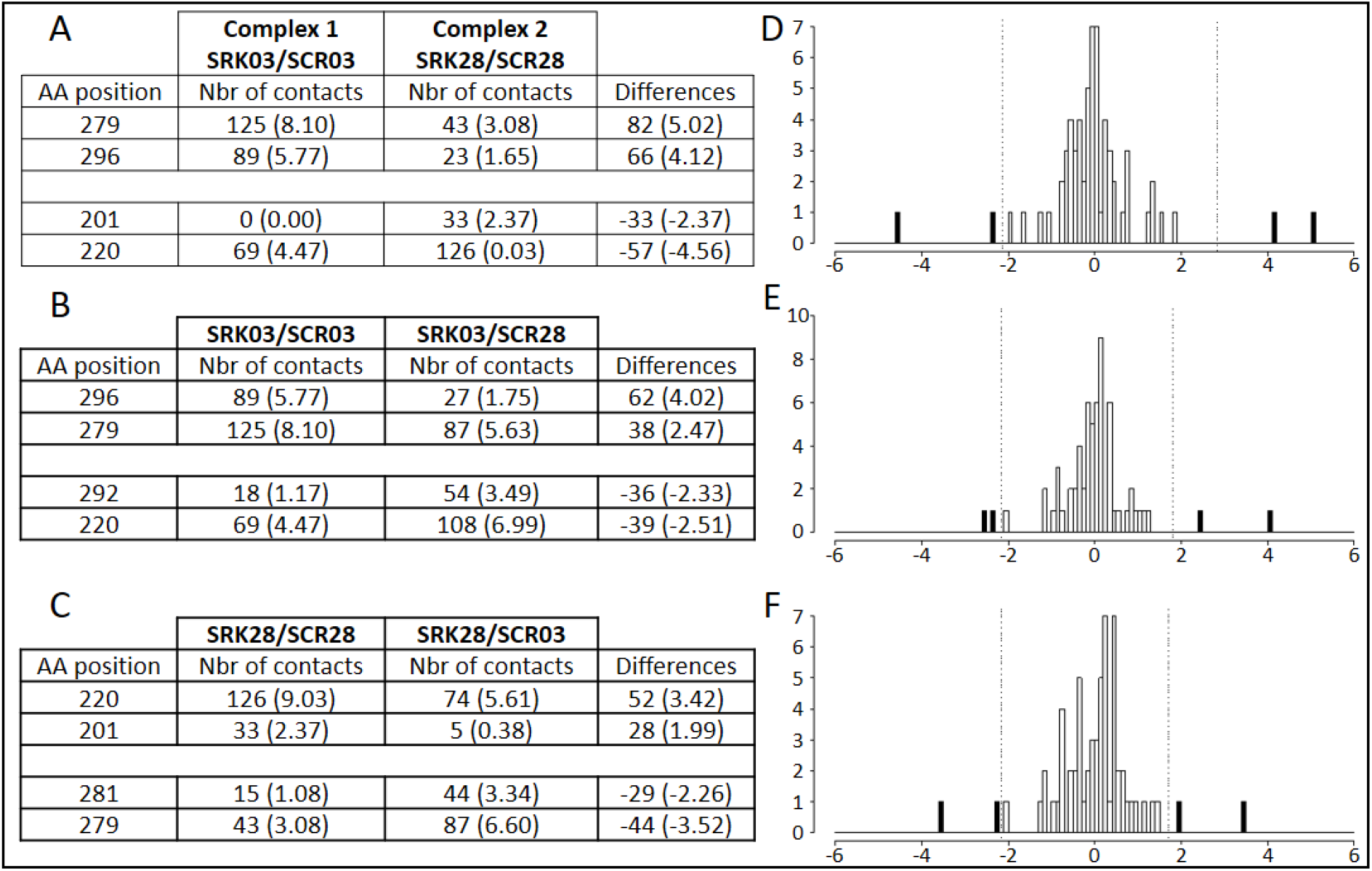
Comparison of amino-acid contacts between the different SCR/SRK complexes points to amino acids of SRK potentially involved in specificity of ligand recognition. The number of atomic contacts between amino-acids of the receptor and the ligand was compared between **(A)** the two cognate complexes (SCR03/SRK03 vs SCR28/SRK28); **(B)** the cognate vs non-cognate complex established by SRK03 (SCR03/SRK03 vs SCR28/SRK03) and **(C)** the cognate vs non-cognate complex established by SRK28 (SCR28/SRK28 vs SCR03/SRK28). The three tables list amino-acid positions who establish the most extreme differences in atomic interactions in each comparison, values in parentheses correspond to the proportion of contacts realized by each aa independently ((number of contact in the complex / number of contact for one aa) × 100). The full distributions are shown in panels **(D), (E)** and **(F)** as the difference of proportion of contacts realized by each aa in both complexes. The vertical lines represent the 5% extreme values. Positive differences correspond to aa involved in numerous contacts in complex 1 compare to complex 2 whereas negative differences correspond to aa involved in numerous contact in complex 2 compare to complex 1.

**Figure S9.**
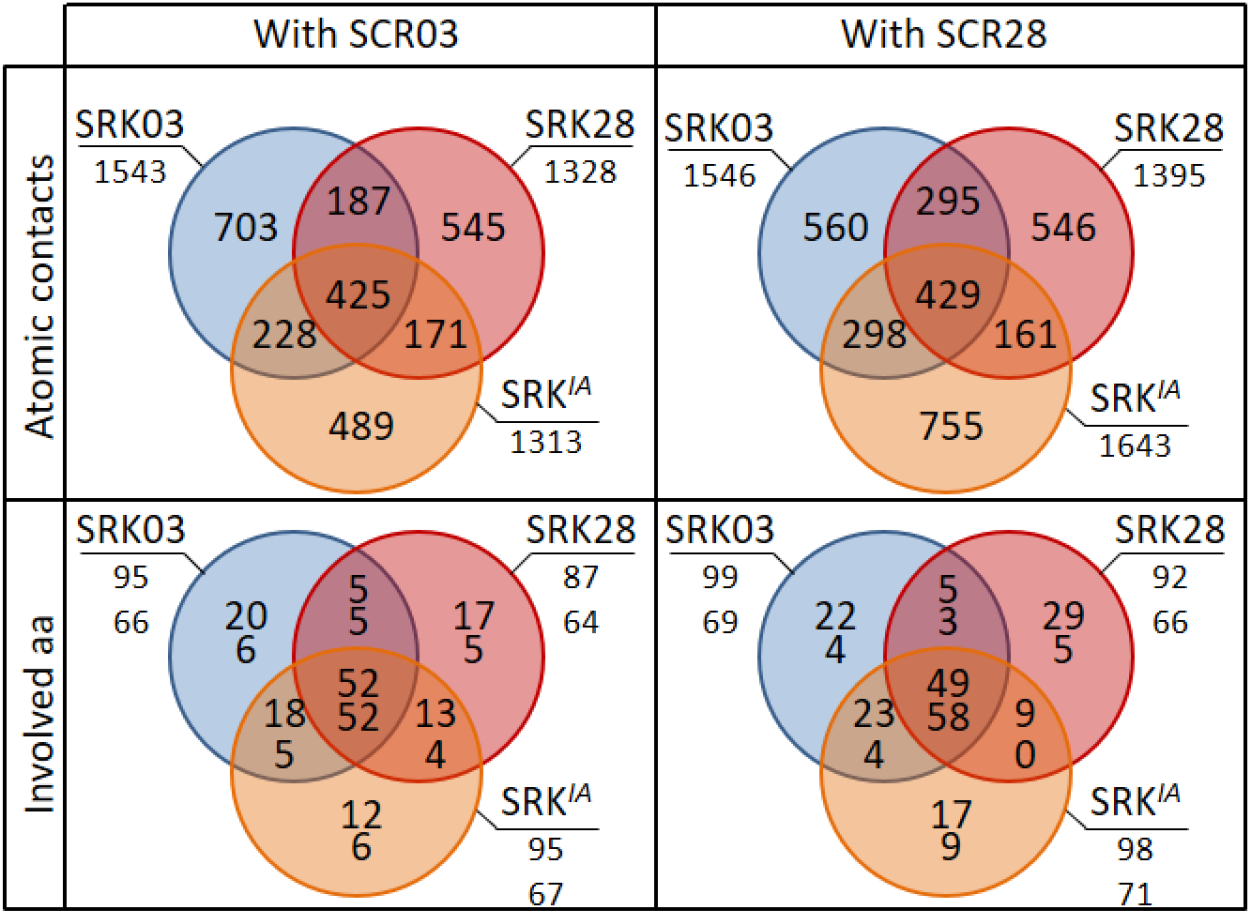
Venn diagram representations of binding features for SRK03, SRK28 and SRKa^IA^ receptors coupled with either SCR03 or SCR28 ligand. Two features were compared: i) Atomic contacts are defined by the nature of the amino acid in both proteins, the chain they belong to, their position and the respective atoms involved in the contact (C, N, O, CA main chain atoms or the side chain atoms Cα, C_β_, C_ε_, …) and ii) The amino acid involved are defined by the nature of the amino acid, the chain they belong to and their position, independently for SRK (numbers on top) and SCR (bottom numbers). For instance, SRKa^IA^ shared 228 atomic contacts with SRK03 and only 171 contacts with SRK28, when complexed with SCR03. This tendency is also apparent when SRKa^IA^ was complexed with SCR28 (298 identical contact with SRK03 but only 161 with SCR28).

**Figure S10:**
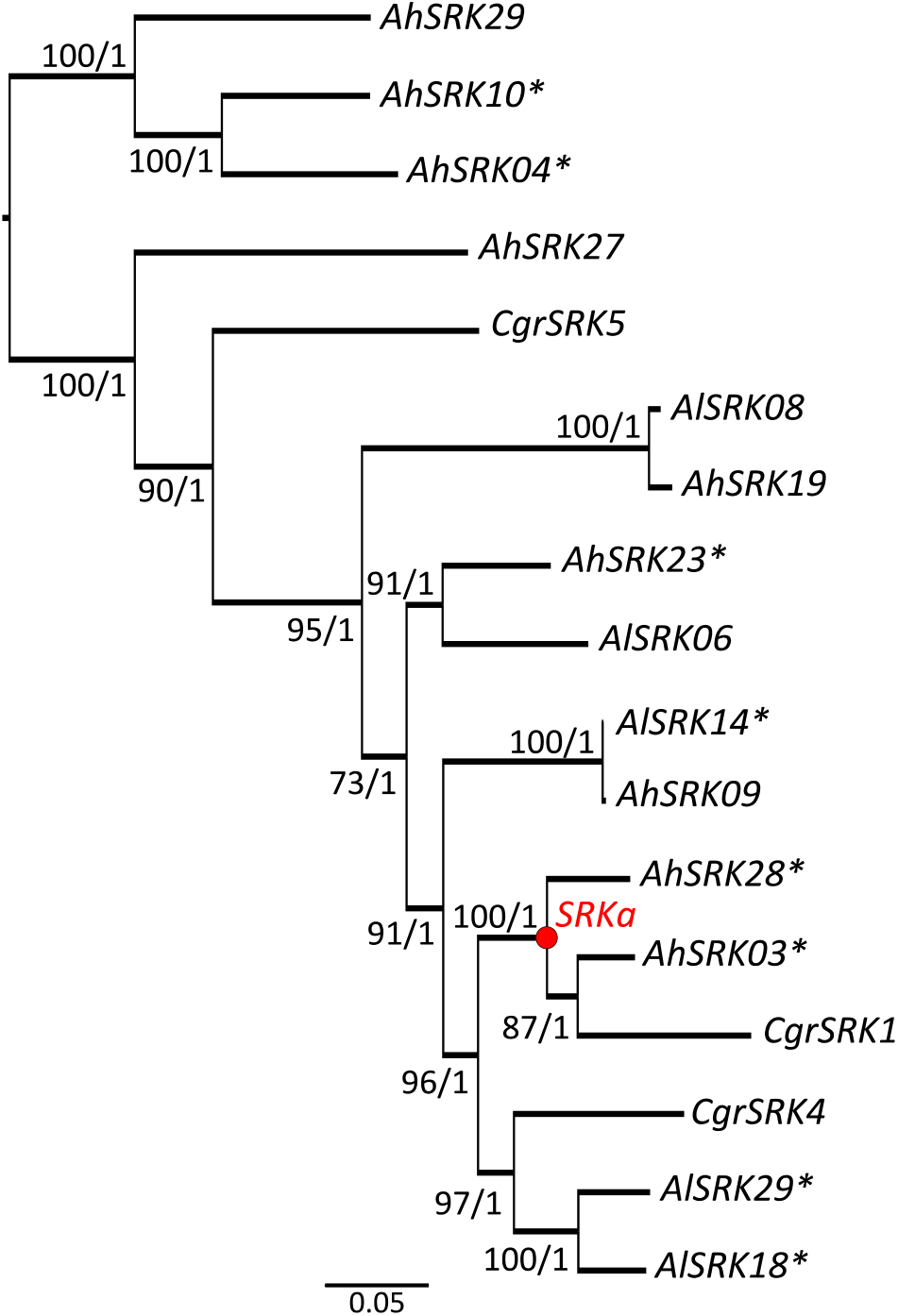
Maximum likelihood phylogenetic tree based on the 17 SRK alleles used for SRKa construction. The Bayesian reconstruction displays an identical topology. Values at the branches represent the bootstrap percentage / posterior probabilities. The node representing the reconstructed ancestral SRK allele (SRKa) is shown in red. Asterisks highlight alleles with a complete sequence available for ancestral reconstruction.

**Figure S11:**
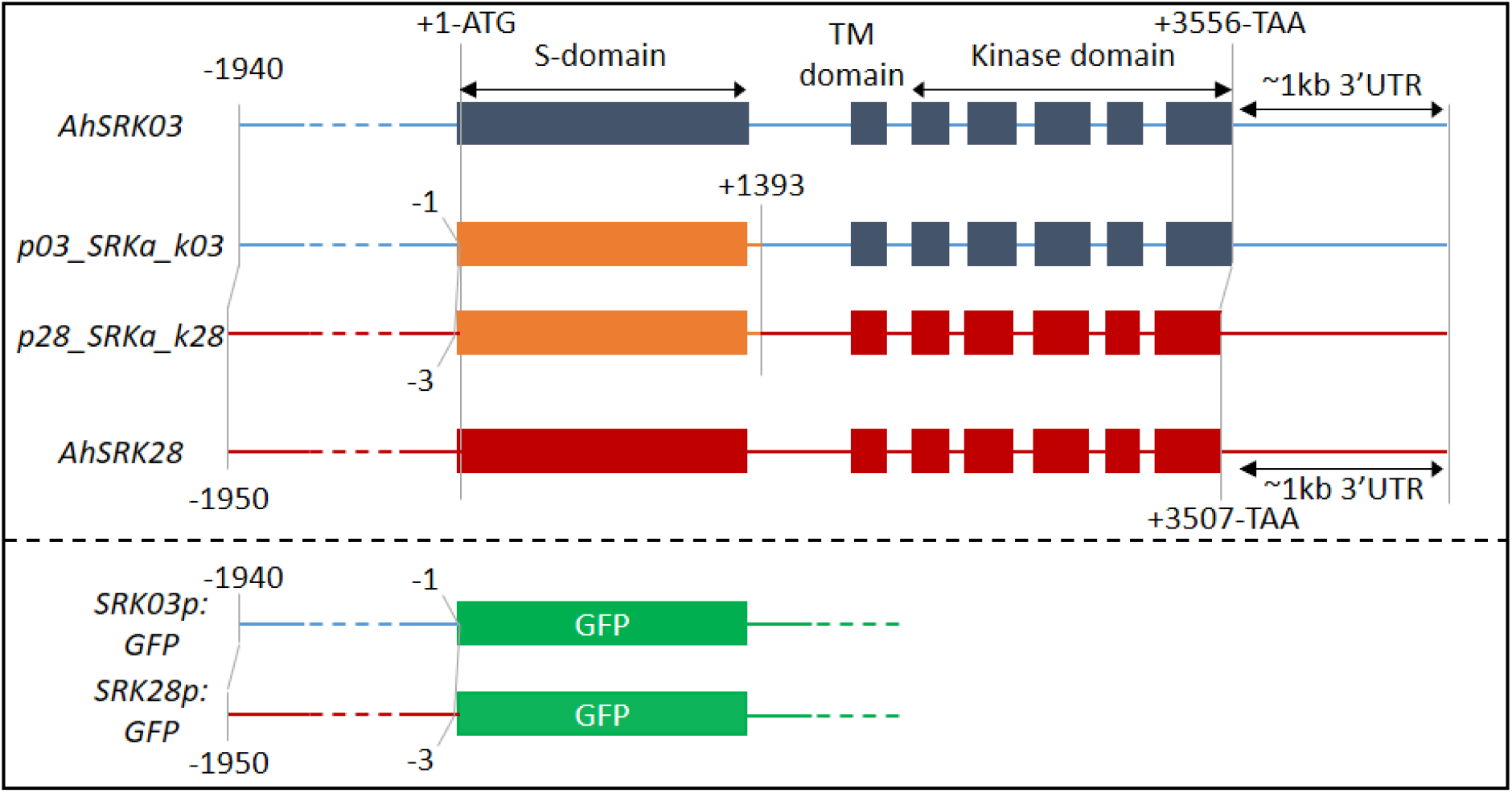
Schematic representation of the molecular constructs used for A. thaliana transformation. AhSRK03 and AhSRK28 constructs were cloned using a unique genomic fragment starting around −2 Kb and ending 1 Kb after the stop codon. SRKa constructs were each cloned using 3 DNA fragments (amplified promoter and kinase domain of AhSRK03 or AhSRK28 and the synthesized ancestral S-domain). The different parts of the construct were concatenated using the Multisite Gateway Technology at the positions indicated on the figure. SRK03p:GFP and SRK28p:GFP constructs were generated using the same promoter sequence as in the SRKa constructs, but were introduced into the pKGWFS7.0 destination vector.

**Table S1:** Detail of the different SRK/SCR interaction protein modelling. Following the eSRK:SCR template structure where two SCR molecules interact with two SRK proteins to form a heterotetramer (Ma et al., 2016), we indicate the two SRK molecules with their chain identifier A and B and the two SCR molecules with G and H. For each complex, the number of involved amino acids and the number of atomic contacts are defined for each protein chain interaction (AG, AH, BG and BH). Underlined numbers in the column “involved aa” correspond to the number of amino acids involved in both cognate and non-cognate interactions.

| SRK receptor | chain | SCR ligand | chain | SRK involved aa | SCR involved aa | Atomic contacts |
| --- | --- | --- | --- | --- | --- | --- |
| S03 | A | S03 | G | <u>16</u>+6=22 | <u>9</u>+9=18 | 464 |
|  | A |  | H | <u>19</u>+13=32 | <u>12</u>+7=19 | 399 |
|  | B |  | G | <u>23</u>+1=24 | <u>7</u>+10=17 | 273 |
|  | B |  | H | <u>15</u>+2=17 | <u>7</u>+9=16 | 407 |
|  |  |  |  |  |  | **1543** |
| S28 | A | S28 | G | <u>15</u>+1=16 | <u>8</u>+7=15 | 277 |
|  | A |  | H | <u>17</u>+7=24 | <u>10</u>+7=17 | 326 |
|  | B |  | G | <u>17</u>+18=35 | <u>4</u>+14=18 | 471 |
|  | B |  | H | <u>15</u>+2=17 | <u>9</u>+7=16 | 321 |
|  |  |  |  |  |  | **1395** |
| S03 | A | S28 | G | <u>16</u>+5=21 | <u>9</u>+9=17 | 366 |
|  | A |  | H | <u>19</u>+8=27 | <u>10</u>+7=17 | 389 |
|  | B |  | G | <u>23</u>+4=28 | <u>4</u>+13=17 | 380 |
|  | B |  | H | <u>15</u>+8=23 | <u>9</u>+9=18 | 411 |
|  |  |  |  |  |  | **1546** |
| S28 | A | S03 | G | <u>15</u>+5=20 | <u>9</u>+6=15 | 378 |
|  | A |  | H | <u>17</u>+10=27 | <u>12</u>+7=19 | 353 |
|  | B |  | G | <u>17</u>+3=20 | <u>7</u>+10=17 | 244 |
|  | B |  | H | <u>15</u>+5=20 | <u>7</u>+10=17 | 353 |
|  |  |  |  |  |  | **1328** |

**Table S2.** Accession numbers for the sequences used in the phylogenetic reconstruction

| Sequence | Accession number |
| --- | --- |
| AhSRK03 | KJ772380.1 |
| AhSRK04 | KJ461484.1 |
| AhSRK08 | EU075130.1 |
| AhSRK09 | EU075131.1 |
| AhSRK10 | KM592810.1 |
| AhSRK19 | EU075140.1 |
| AhSRK23 | EU878008.1 |
| AhSRK27 | EU878012.1 |
| AhSRK28 | KJ461478.1 |
| AhSRK29 | KM592798.1 |
| AlSRK06 | GQ351354.1 |
| AlSRK08 | JX464638.1 |
| AlSRK14 | KJ772405.1 |
| AlSRK18 | KJ772412.1 |
| AlSRK29 | AY186776.1 |
| AlSRK39 | KJ772418.1 |
| CgrSRK1 | DQ530637.1 |
| CgrSRK4 | DQ530640.1 |
| CgrSRK5 | DQ530641.1 |
| CgrSRK6 | DQ530642.1 |

**Table S3.**
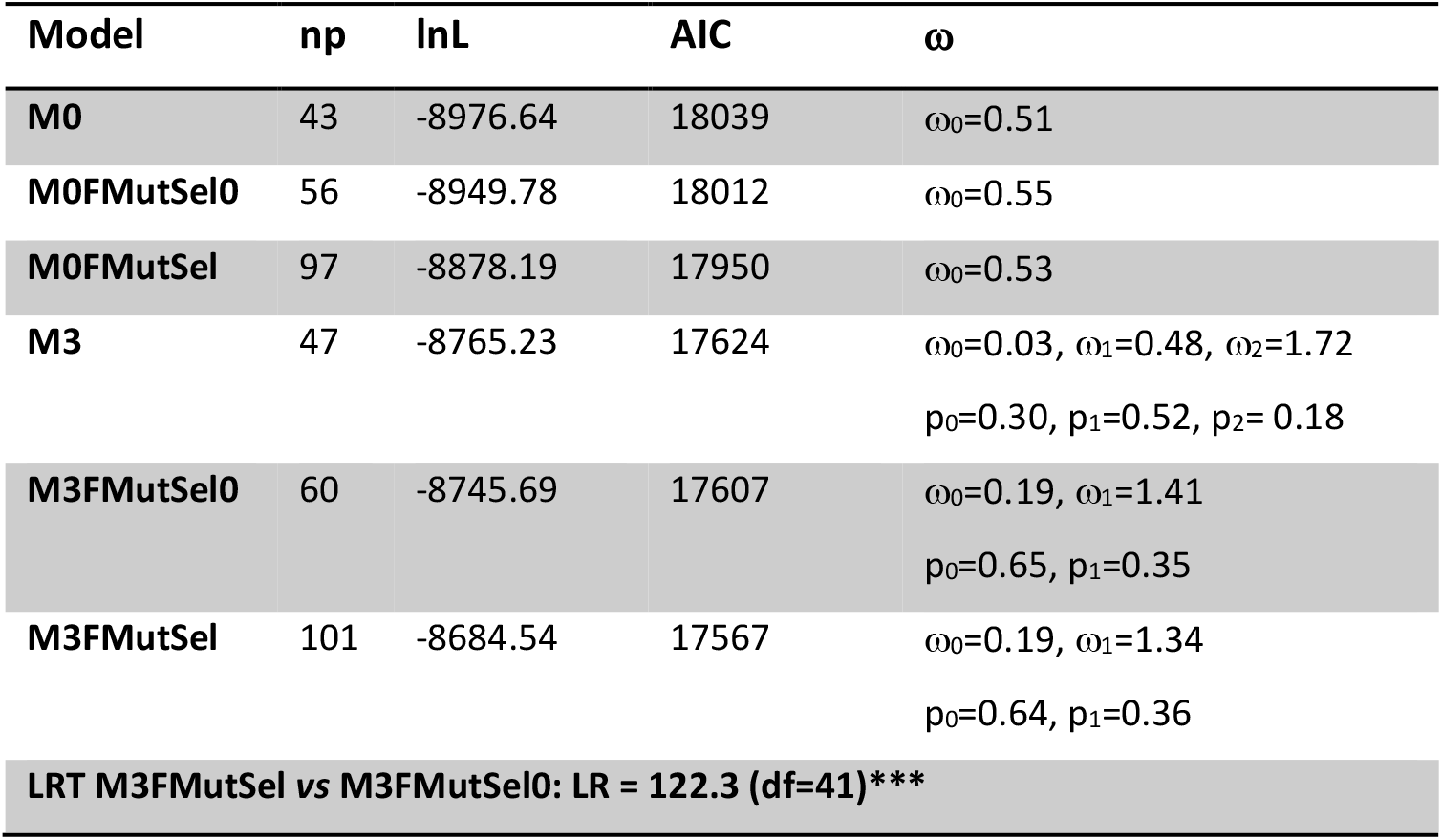
PAML ancestral analyses of the SRK protein and model comparison. np is the number of parameters in the model; lnL is the log likelihood score; AIC (Akaike Information criterion = −2*lnL+2*np) is a measure of the goodness of fit of an estimated statistical model; ω□is the nonsynonimous/synonymous substitution ratio; LR is the likelihood ratio: df is the degree of freedom in LRT (Likelihood Ratio Test); *** Highly significant (P-value < 0.0001).

| Model | np | lnL | AIC | $\omega$ |
| --- | --- | --- | --- | --- |
| M0 | 43 | -8976.64 | 18039 | $\omega_0=0.51$ |
| M0FMutSel0 | 56 | -8949.78 | 18012 | $\omega_0=0.55$ |
| M0FMutSel | 97 | -8878.19 | 17950 | $\omega_0=0.53$ |
| M3 | 47 | -8765.23 | 17624 | $\omega_0=0.03, \omega_1=0.48, \omega_2=1.72$<br>$p_0=0.30, p_1=0.52, p_2=0.18$ |
| M3FMutSel0 | 60 | -8745.69 | 17607 | $\omega_0=0.19, \omega_1=1.41$<br>$p_0=0.65, p_1=0.35$ |
| M3FMutSel | 101 | -8684.54 | 17567 | $\omega_0=0.19, \omega_1=1.34$<br>$p_0=0.64, p_1=0.36$ |
| LRT M3FMutSel vs M3FMutSel0: LR = 122.3 (df=41)*** |  |  |  |  |

**Table S4:**
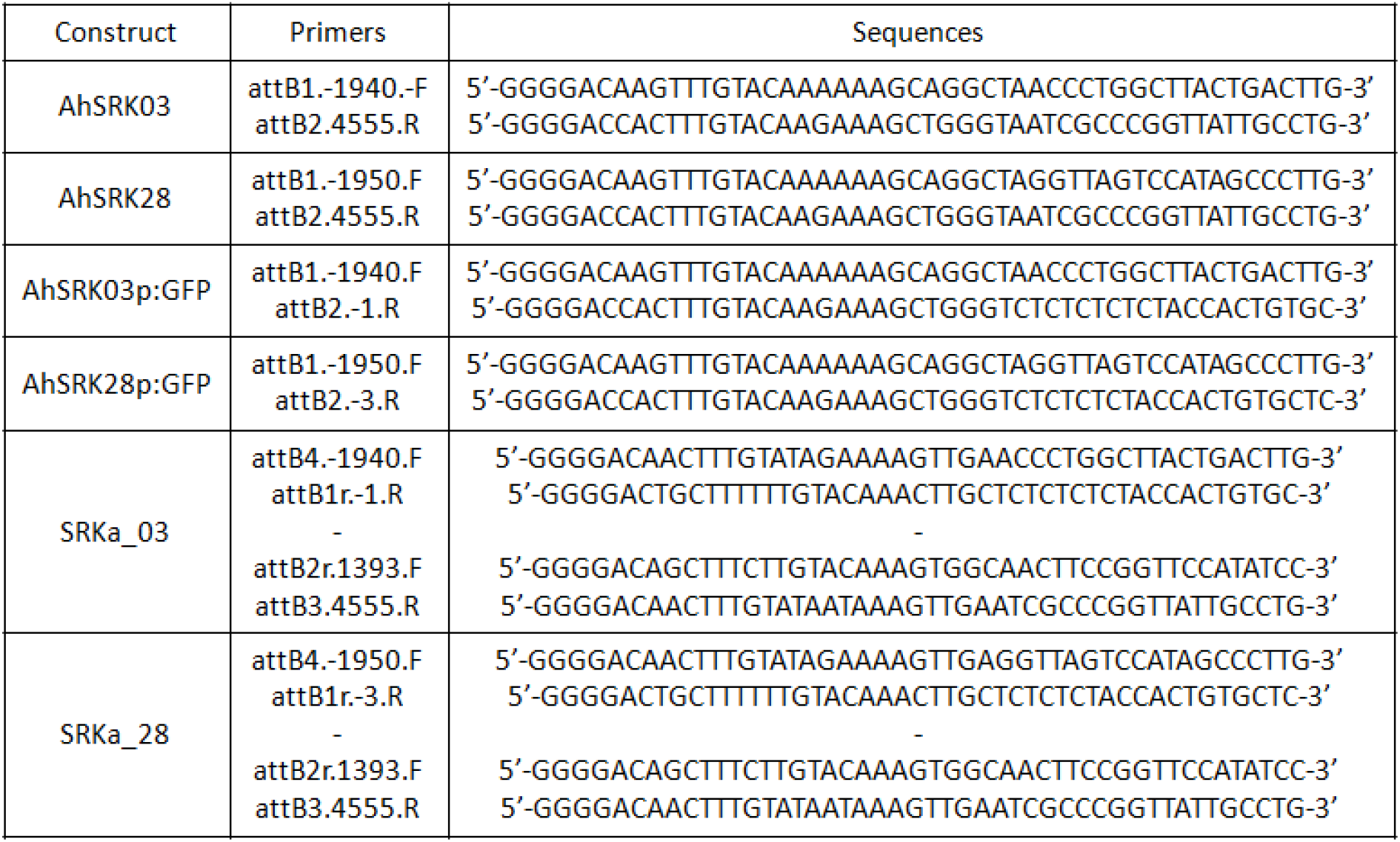
Gateway primers used for molecular constructs.

